# DOT1L activity affects cell lineage progression in the developing brain by controlling metabolic programs

**DOI:** 10.1101/2022.04.08.487591

**Authors:** Bismark Appiah, Camilla L. Fullio, Christiane Haffner, Patrice Zeis, Martin Treppner, Patrick Bovio, Arquimedes Cheffer, Ilaria Bertani, Harald Binder, Dominic Grün, Nereo Kalebic, Elena Taverna, Tanja Vogel

## Abstract

Cortical neurogenesis depends on the tight balance between self-renewal and differentiation of apical progenitors (APs), the key progenitor type generating all other neural cells including neocortical neurons. We here report the activity of the histone methyltransferase DOT1L as a gatekeeper for AP cell identity. Combining lineage tracing with single-cell RNA sequencing of clonally related cells, we explore consequences of DOT1L inhibition on AP lineage progression during neurogenesis in the embryonic mouse neocortex. At the cellular level, DOT1L inhibition led to increased neurogenesis driven by a shift from asymmetric self-renewing to symmetric neurogenic divisions of APs. At the molecular level, we show that DOT1L activity preserved AP identity by promoting transcription of a gene set involved in AP metabolism. On a mechanistic level, DOT1L inhibition increased expression of metabolic genes, including microcephaly-associated Asparagine synthetase (*Asns*) and overexpression of ASNS in APs resulted in increased neuronal differentiation. *Asns* expression was predicted to be controlled through EZH2 and we show that DOT1L activity allows PRC2-mediated repression of *Asns* expression. Importantly, inhibition of ASNS activity rescued increased AP differentiation upon DOT1L inhibition. Our data show that DOT1L activity/PRC2 crosstalk controls AP lineage progression by regulating AP metabolism, and they provide a mechanistic view on how DOT1L activity might affect neocortical neurogenesis.

## Introduction

The six-layered cerebral cortex has been studied extensively with regard to the transcription factors (TFs) governing the determination of cell identity of stem and progenitor cells, as well as the generation of neurons during embryonic development and in neurodevelopmental disorders (NDD) (Molyneaux et al., 2007; Arnold et al., 2008; Britanova et al., 2008; Bedogni et al., 2010; Clark et al., 2020). Neural stem and progenitor cells are classified into two main types based on the location of mitosis: apical progenitors (APs) that divide at the apical surface of the ventricular zone, and basal progenitors (BPs) that typically divide in the subventricular zone (SVZ) (Taverna et al., 2014). In rodents, APs and BPs differ not only in localisation but also in their mode of division and neurogenic potential. At mid-neurogenesis mouse APs mainly undergo asymmetric, self-renewing division to generate one AP and one BP. In contrast, mouse BPs are mainly undergoing symmetric neurogenic consumptive division to generate two neurons (Taverna et al., 2014).

The relevance of maintaining a tight balance between different modes of divisions is illustrated by primary microcephaly (Fish et al., 2006), a human NDD, where the premature switch to neurogenic consumptive division was identified as a key driver of the dramatic reduction in brain size observed in patients. Recent work has identified TF activities, mitotic spindle components (Haydar et al., 2003) and specific metabolic signatures (Lange et al., 2016; Zheng et al., 2016; Journiac et al., 2020) as potential molecular regulators of the AP division mode. Among TFs, master regulators of AP identity such as SOX2, PAX6 and EMX2 were reported to promote the symmetric divisions of APs (Heins et al., 2001; Estivill-Torrus et al., 2002; Asami et al., 2011; Hagey and Muhr, 2014).

But despite the progress in the definition of the genetic logic of cell fate determination and acquisition of cell identity (Molyneaux et al., 2007), little is still known about the epigenetic regulation of the division mode, an aspect that is crucially linked to the symmetry and asymmetry of cell fate specification in APs. Defining the relationship between epigenetics and symmetry/asymmetry of division is important considering that (i) epigenetic information is heritable and can affect cell fate decision (Hirabayashi and Gotoh, 2010) and (ii) histone proteins can be asymmetrically partitioned during cell division (Wooten et al., 2020; Roubinet et al., 2021). These data demand for extending the sparse insights as of yet into the cellular and mechanistic link between epigenetic regulation and NDDs (Bjornsson, 2015; Mastrototaro et al., 2017).

We recently reported that the chromatin modifier Disruptor of telomeric silencing-like 1 (DOT1L) affects cortical development. Genetic inactivation of DOT1L at early stages of neurogenesis (around E9.5 in mice) depleted the progenitor pool and resulted in premature differentiation to neurons (Franz et al., 2019). Moreover, pharmacological inhibition of DOT1L in cultured mouse neural progenitor cells (NPCs) led to increased neuronal differentiation, which correlated with DOT1L-mediated alteration of accessibility of SOX2-bound enhancer regions (Ferrari et al., 2020). Interestingly, DOT1L is linked to genes known to regulate the symmetry and asymmetry of cell division and cell metabolism in neural stem and progenitor cells (Roidl et al., 2016; Franz et al., 2019). These observations prompted us to study how DOT1L affects specifically AP behaviour and lineage progression in the cerebral cortex *in vivo* during mid-neurogenesis (around E14.5 in mice).

By using single-cell labelling and two lineage tracing techniques, in combination with single-cell RNA sequencing (scRNA-seq) and cell biological enquiry, we show that DOT1L affects the division mode, the choice between self-renewal and differentiation, and ultimately the maintenance of AP identity during cortical development. Mechanistically, our data show that DOT1L activity maintains AP self-renewal potential, by hampering the premature activation of metabolic programs of differentiated cells, and by preventing dissipation of the poised promoter state and conserving PRC2-mediated H3K27me3 silencing.

## Results

### DOT1L inhibition increases delamination

As previous data suggest that DOT1L controls the generation of neurons, we sought to understand the underlying cellular mechanism by pharmacologically blocking DOT1L activity in the mouse developing brain during mid-neurogenesis. To this end, we treated mouse E14.5 hemispheres with EPZ5676 (EPZ), a widely used inhibitor of DOT1L in cancer research (Nassa et al., 2019; Vlaming et al., 2019; Vatapalli et al., 2020) and during reprogramming (Cao et al., 2018), for 24 h in the hemisphere rotation (HeRo) culture (Schenk et al., 2009) (**Fig. 1A**). In the rodent neocortex, the acquisition of a basal fate (BP or neuron) is concomitant with the relocation of the centrosome and associated cilia from the ventricular surface to an abventricular location. Using immunofluorescence for organelle-specific markers, we assessed the number and distribution of centrosomes (as revealed by TUBG1 staining, **Fig. 1B**) and primary cilia (as revealed by ARL13B staining, **Fig. 1B**) in the cortical wall as a proxy for delamination (Taverna et al., 2011; Wilsch-Brauninger et al., 2012; Tavano et al., 2018). When assessing the entire cortical wall, these parameters were not affected upon EPZ treatment (**Fig. 1C, D**). However, when scoring the distribution of TUBG1- and ARL13B-positive structures specifically in the VZ, we detected a strong increase in abventricular TUBG1- and ARL13B-positive structures in EPZ-treated samples compared to control along with a parallel decrease in ventricular centrosomes and cilia (**Fig. 1E-G**). These data strongly suggest that DOT1L inhibition increases AP delamination, possibly prompting a basal cell fate.

**Figure 1.**
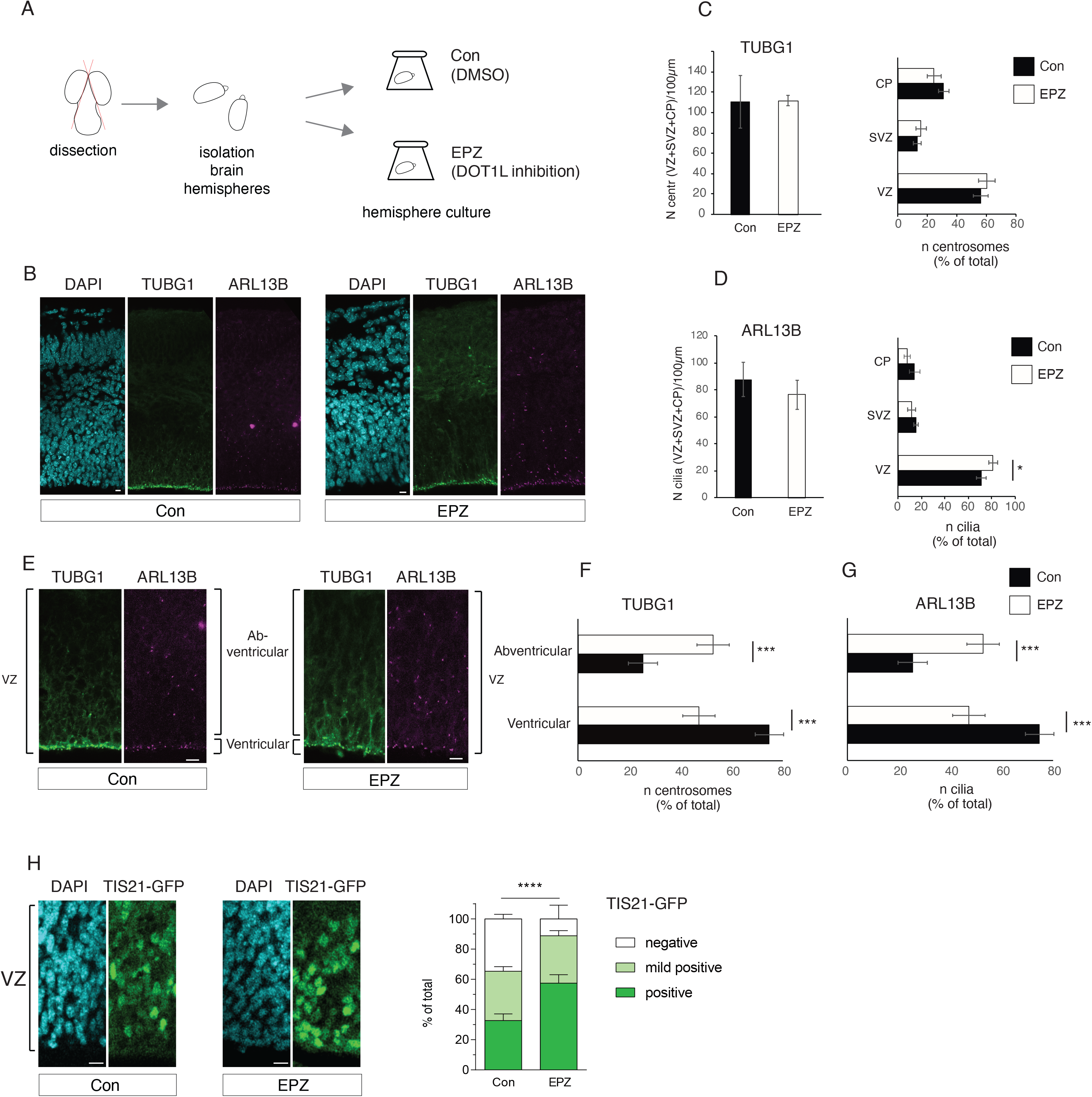
Basal localization of centrosomes and cilia in the VZ indicate delamination of progenitors upon DOT1L inhibition. **A)** Outline of experiment. **B)** Overview of immunofluorescence staining for centrosomes (TUBG1) and cilia (ARL13B) in control (Con) (left) and EPZ treated sections (right). **C)** Left: quantification of total centrosome numbers in control and EPZ treated sections in 100 µm. Right: number of centrosomes in ventricular zone (VZ), sub-ventricular zone (SVZ), and cortical plate (CP) expressed as percentage of total per condition. **D)** Left: quantification of total cilia numbers in control and EPZ treated sections in 100 µm. Right: number of cilia in VZ, SVZ, and CP expressed as percentage of total per condition. **E)** Overview of VZ showing TUBG1 and ARL13B immunofluorescence staining. Squared parentheses indicate the ventricular and abventricular compartment. **F)** Quantification of TUBG1- and **G)** ARL13B-positive structures in ventricular and abventricular compartment expressed as percentage of the total number per condition. All distribution of TUBG1 and ARL13B are based on quantifications of 4 fields of view including the relevant area to quantify (VZ, SVZ, CP), from 4 independent experiments (n=4; for the distribution in the cortical wall: Con: 755 ARL13B-positive structures and 1123 TUBG1-positive structures; EPZ: 715 ARL13B- positive structures and 1123 TUBG1-positive structures; for the distribution in the VZ, Con: 633 ARL13B-positive structures and 1005 TUBG1-positive structures; EPZ: 686 ARL13B-positive structures and 1023 TUBG1-positive structures). T-test, two-tails was performed for all quantifications, *p<0.05, ***p<0.001, error bars represent SD. **H)** Left: Immunofluorescence staining showing overview of TIS21-GFP-positive (GFP+) cells in VZ of control and EPZ treated sections. Right: Graph comparing total GFP+ cells in VZ for control and EPZ conditions. Chi-square test was performed, ****p<0.0001, error bars represent SEM. Scale bar in all panels: 10 µm.

We next used *ex utero* electroporation of a mCherry reporter and subsequent HeRo culture for 24 h in presence or absence of EPZ to assess DOT1L effects on cell positioning (**Supplementary Fig. 1A**). Whereas the control mCherry-positive cell bodies resided within the apical-most half of the VZ, upon EPZ treatment mCherry-positive cell bodies localised more basally (**Supplementary Fig. 1B**).

To assess the effect of DOT1L inhibition on the neurogenic or proliferative potential of APs, we used the *Tis21(Btg2)-Gfp* mouse, where *Gfp* is under the control of the promoter of *Tis21* (*Btg2*), a neurogenic marker (Haubensak et al., 2004). This mouse model allows following neurogenesis by monitoring the appearance and localization of GFP (**Fig 1H**, left panels). Upon DOT1L inhibition we observed an increase in the proportion of cells positive for TIS21-GFP (**Fig. 1H**), suggesting that the acquisition of a basal fate was paralleled by a switch to neurogenesis (Haubensak et al., 2004). Our data combining cell biological enquiry, histological analysis of cell distribution and marker expression, suggest that upon DOT1L inhibition the AP progeny is committed to a basal and differentiated fate.

### DOT1L inhibition promotes neuronal differentiation

To dissect the underlying cellular events at a higher temporal and spatial resolution, we used manual microinjection for targeting single AP in tissue (Taverna et al., 2012; Wong et al., 2014). APs were microinjected with Dextran-Alexa555 in organotypic slices from E14.5 mouse telencephalon (see scheme in **Fig. 2A, Supplementary Fig. 2A**). The slices were kept in culture for various time windows to reconstruct and study the AP progeny. Cell identity was defined by combining several criteria, such as cell morphology, location and marker expression (Taverna et al., 2011; Wong et al., 2014; Kalebic et al., 2016; Tavano et al., 2018; Shull et al., 2019). 3 h after microinjection the progeny of microinjected APs resided almost exclusively in the VZ and had a contact with the ventricle (**Supplementary Fig. 2B**). Instead, 24 h after microinjection the progeny of microinjected APs (i) located not only to the VZ, but also to the SVZ and CP, and (ii) expressed the BP marker EOMES and the neuronal marker TUBB3 (**Supplementary Fig. 2C**, panels on the right).

**Figure 2.**
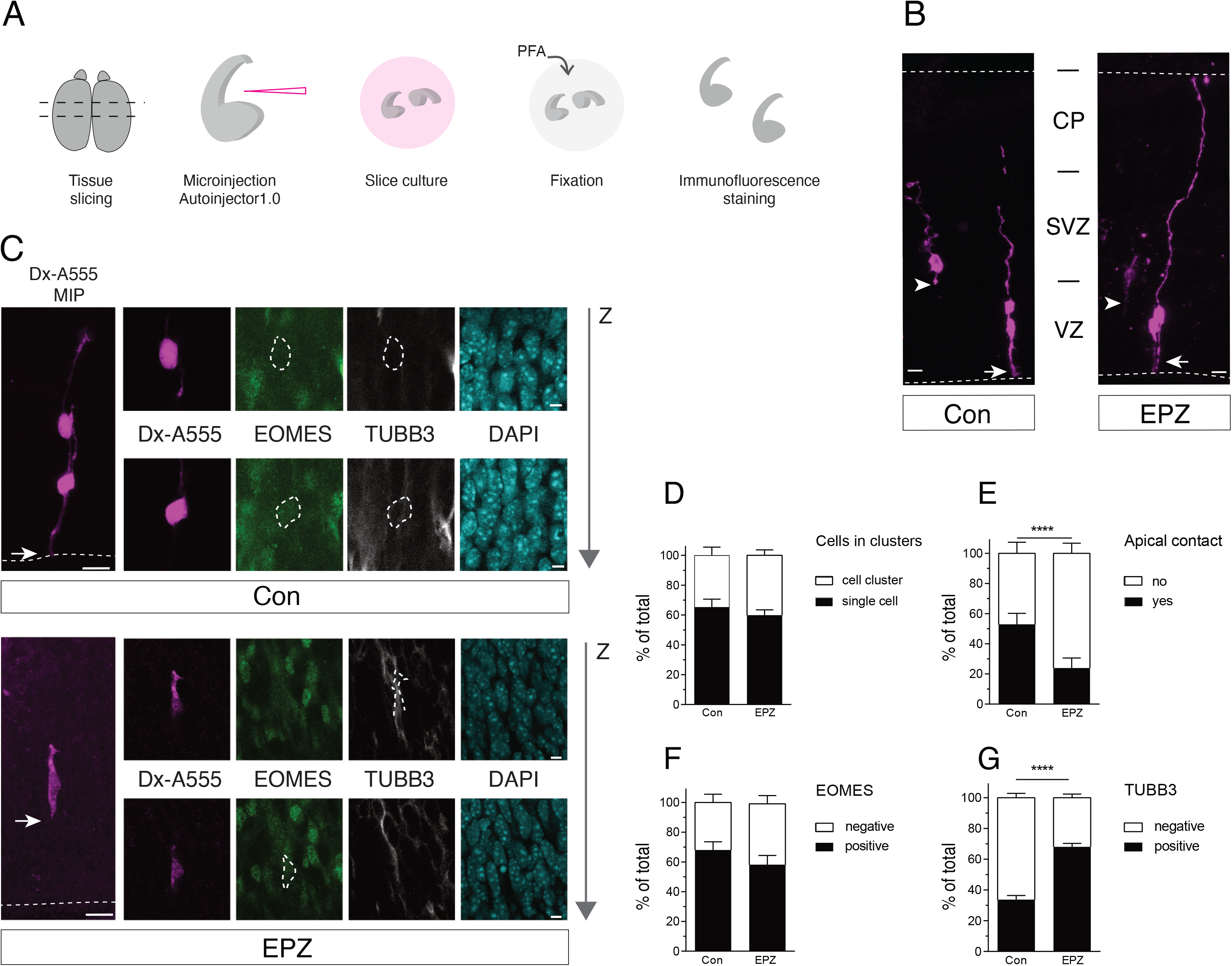
Lineage tracing of microinjected cortical APs shows DOT1L inhibition increases neurogenesis and apical delamination. **A)** Outline of experiment. **B)** Microinjected section treated with DMSO (Con) or EPZ showing Dx-A555 labelled cells after 24 h in slice culture. White arrow indicates the apical process contacting the ventricular surface. White arrowheads indicate the apical-directed process of delaminated cells. **C)** Immunofluorescence staining of microinjected sections showing the progeny of microinjected cells (a daughter cell pair) expressing EOMES and TUBB3. White arrows indicate the apical-directed processes. Note that in the top panel the apical-directed process contacts the ventricular surface (dotted white line), while in the bottom panel it does not contact the ventricular surface, indicating that the cell has delaminated. **D)** Quantification of microinjected cells found as single-cell or two-cells clusters, expressed as percentage of total Dx-A555-positive cells per condition. **E)** Quantification of microinjected cells with or without contact to the apical surface of the organotypic slice expressed as percentage of total Dx-A555-positive cells per condition. Total n of independent microinjection experiments: 6 (n=3 wild type embryos, n=3 in Tis21-GFP embryos). Total number of cells scored: Con, 156 cells; EPZ, 287 cells. **F)** EOMES+ and EOMES- cells plotted as a percentage of total cells scored per condition. **G)** TUBB3-positive (TUBB3+) and TUBB3-negative (TUBB3-) cells in control and EPZ conditions plotted as a percentage of total cells scored per condition. Cell scoring based on TUBB3 or EOMES expression are from three independent experiments (n=3; Con: 70 cells, EPZ: 113 cells). Fisher’s exact test was performed for all quantifications, ****p<0.0001, error bars represent SEM. MIP - maximum intensity projection. VZ - Ventricular Zone, SVZ - Subventricular Zone, CP - Cortical Plate. Scale bars in all main panels 20 µm, and all insets 5 µm.

Having hence successfully implemented AP microinjection, we applied it to study at the single-cell level the effect of DOT1L inhibition on cell morphology, cell identity and lineage progression (**Fig. 2B-G**). We first analysed the general composition of the progeny of microinjected cells and found that EPZ treatment did not change the proportion of cells found in two-cell clusters as opposed to single-cells (**Fig. 2C, D**). These data suggest that EPZ does not have unspecific effects on cell cycle progression *per se*, nor that it causes mitotic arrest. The morphological analysis showed that the inhibition of DOT1L resulted in an increase in the number of cells without apical contact (**Fig. 2B, C, E**), suggesting the generation of a delaminated and more differentiated cell progeny (either BPs or neurons) compared to APs. Double staining with EOMES and TUBB3 revealed a slight decrease in the fraction of EOMES-expressing cells (BPs; **Fig. 2C, F**) compared to a strong increase in the proportion of TUBB3-expressing cells (neurons; **Fig. 2C, G**). Taken together, these data suggest that the inhibition of DOT1L might favour the acquisition of a neuronal over BP cell fate.

### DOT1L inhibition promotes symmetric neurogenic division

To follow the switch to neurogenesis at single-cell resolution, microinjection into single-cells was performed in the *Tis21(Btg2)-Gfp* mouse. EPZ treatment of the respective slice cultures increased the proportion of microinjected cells that are TIS21- GFP positive in the neocortex (**Fig. 3A, B**).

**Figure 3.**
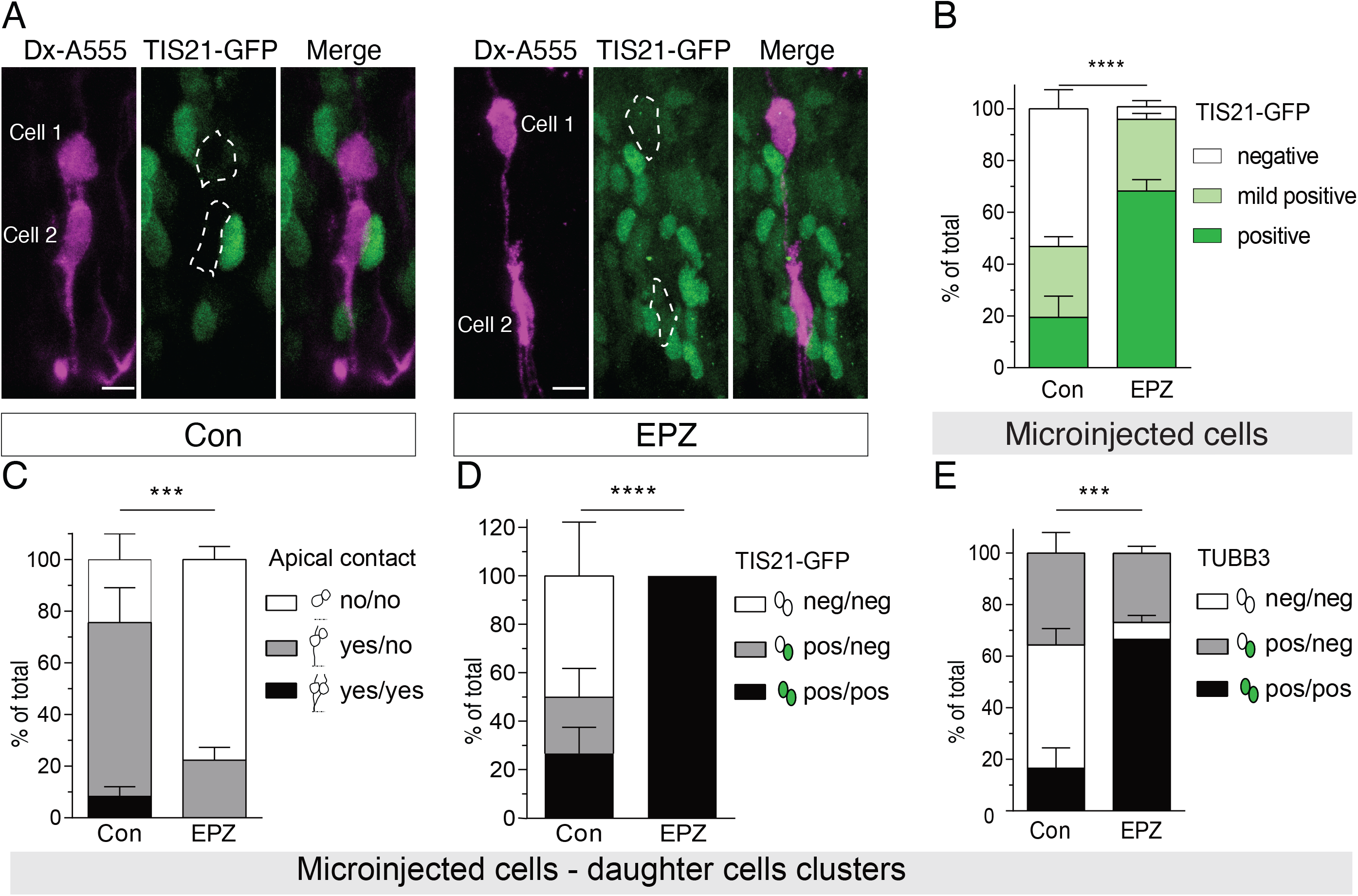
DOT1L inhibition leads to neurogenic commitment of VZ progenitors. **A)** Microinjected cells in the VZ (magenta) of control (Con) and EPZ treated slices 24 h after microinjection and slice culture. Left: daughter cell pair in control section with asymmetric GFP expression, with cell 1 (basal location) being GFP-positive, and cell 2 (apical location) GFP-negative. Right: EPZ treated section showing a daughter cell pair with symmetric GFP+ expression. **B)** Graph showing proportion of labelled cells expressing GFP+, GFP mild+, or GFP- expressed as percentage of total cells per condition. **C)** Symmetry or asymmetry in daughter cells’ apical contact (+/+: both cells have apical contact; +/-: one cell has apical contact and the other not; -/-: both cells lack apical contact). **D)** Symmetry (GFP+/+, GFP-/-) or asymmetry (GFP+/-) in GFP expression in daughter cells expressed as percentage of total cells per condition. **E)** Symmetry or asymmetry in TUBB3 expression in daughter cells plotted (+/+: both cells express TUBB3; +/-: one cell expresses TUBB3 and the other not; -/-: none of the cells expresses TUBB3). For quantifications the following number of cells were scored: TUBB3 - 52 daughter cell pairs in total, Con: 19, EPZ: 33 pairs; EOMES - 43 daughter cell pairs in total, Con: 16, EPZ: 27 pairs; TIS21-GFP - 64 daughter cell pairs in total, Con: 35, EPZ: 29 pairs; apical contact - 56 daughter cell pairs in total, Con: 19, EPZ: 37 pairs. Quantifications are from three independent experiments (n=3), 260 cells in total (Con: 86 cells, EPZ: 174 cells). Chi-square test was performed for all quantifications, error bars represent SEM, ***p<0.001, ****p<0.0001, alpha = 0.05. MIP - maximum intensity projections. VZ - ventricular zone. Scale bars: 10 µm.

To address the effects of DOT1L inhibition on the symmetry and asymmetry of AP cell division, we also made use of the high single-cell resolution provided by microinjection. Here, we analysed specifically two-daughter cell clusters, and assessed their symmetry/asymmetry in terms of morphology and cell identity (**Fig. 3C-E**). Morphological analysis revealed that EPZ treatment increased specifically the proportion of two-cell clusters where both cells delaminated (**Fig. 3C**), suggesting that the increased delamination (shown also in **Fig. 2**) derived from a switch to symmetric division, rather than direct delamination of APs.

The analysis of TIS21-GFP in two-daughter cells clusters revealed that upon EPZ treatment both daughter cells expressed TIS21-GFP (**Fig. 3D**). Moreover, in most clusters both daughter cells expressed (symmetrically) the neuronal marker TUBB3 (**Fig. 3E**). Taken together, these data strongly suggest that DOT1L inhibition changes the mode of division of APs from asymmetric self-renewing to symmetric neurogenic division.

### DOT1L inhibition alters the composition of the progenitor populations and favours neurogenesis

To gain insight on the transcriptional signature of the underlying cellular alterations upon DOT1L inhibition, we performed scRNA-seq, focusing exclusively on lineage related cells. To this goal, we specifically labelled single APs before the start of the pharmacological treatment, using two different approaches: (i) microinjection of fluorescent dye (either Dextran-Alexa488 or Dextran-Alexa555) into single neural stem cells using a recently developed robotic microinjection system (Shull et al., 2019) (scheme in **Fig. 4A**, left; **Fig. 4B-E**), and (ii) *ex utero* hemisphere electroporation of a GFP-expressing construct (Calegari et al., 2002) (scheme in **Fig. 4A**, right; **Fig. 4F-H**) (Schenk et al., 2009; Kalebic et al., 2019). In both cases the tissue (tissue slices for microinjection or hemispheres for *ex utero* electroporation) was kept in culture for 24 h and the labelled cells were recovered by FACS (microinjection into single neural stem cells: 307 cells (Con) and 439 cells (EPZ); *ex utero* electroporation: 371 cells (Con) and 379 cells (EPZ)). With both labelling approaches, the number and the quality of cells recovered were sufficient for subsequent scRNA-seq analysis (**Supplementary Fig. 3**).

**Figure 4.**
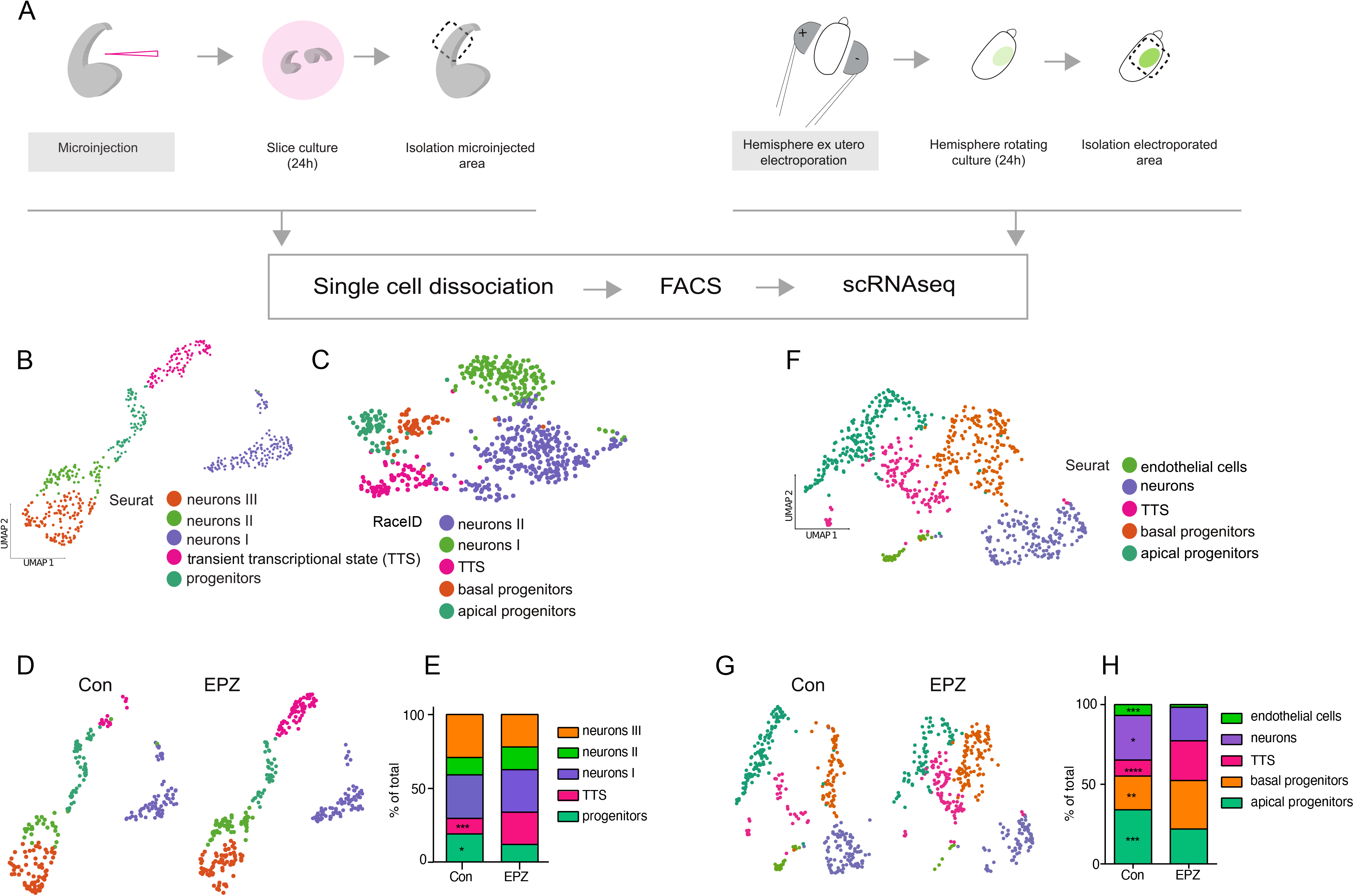
Single-cell transcriptome 24 h after AP labelling via microinjection or *ex utero* electroporation reveals increased neuronal differentiation upon DOT1L inhibition. **A**) Outline of the experiments. **B**) Two-dimensional representation (UMAP embedding) of clusters arising from Seurat analysis of scRNA-seq data from cells labelled via microinjection. A total of 589 cells were retrieved. **C**) Clusters retrieved via RaceID analysis of scRNA-seq data from a total of 639 cells labelled via microinjection represented in low dimension (t-SNE). Cells from both Con and EPZ conditions were combined for clustering analysis. Each dot represents a cell, and cells marked by a colour represent a specific cell state/type. Annotated cells (RaceID): apical progenitors (Con: 34 cells, EPZ: 33 cells), cells in TTS (Con: 26 cells, EPZ: 57 cells), basal progenitors (Con: 29 cells, EPZ: 24 cells), neurons I (Con: 58 cells, EPZ: 92 cells), neurons II (Con: 117 cells, EPZ: 169 cells). **D**) scRNA-seq clusters from microinjected cells split by condition (Con: 248 cells, EPZ: 341 cells). Annotated cells from microinjection data (Seurat v3): apical/basal progenitors (Con: 48 cells, EPZ: 42 cells), cells in transient transcriptional state (TTS) (Con: 26 cells, EPZ: 74 cells), neurons I (Con: 73 cells, EPZ: 98 cells), neurons II (Con: 29 cells, EPZ: 52 cells), neurons III (Con: 72 cells, EPZ: 75 cells). **E**) Fraction of cell states in microinjected cells expressed as percentage of total cells per condition. Fisher’s exact test was performed. *p<0.05, ***p<00.1. **F**) Low dimensional representation (UMAP embedding) of clusters arising from Seurat analysis of scRNA-seq data from cells labelled via electroporation. A total of 711 cells were retrieved. **G**) scRNA-seq clusters from electroporated cells split by condition (Con: 350 cells, EPZ: 361 cells). Annotated cells from electroporation data (Seurat v3): apical progenitors (Con: 119 cells, EPZ: 79 cells), cells in TTS (Con: 35 cells, EPZ: 90 cells), basal progenitors (Con: 74 cells, EPZ: 110 cells), neurons (Con: 98 cells, EPZ: 76 cells), endothelial cells (Con: 24 cells, EPZ: 6 cells). **H**) Fraction of cell states in electroporated cells expressed as percentage of total cells per condition. Fisher’s exact test was performed. *p<0.05, **p<0.01, ***p<0.001, ****p<0.0001.

We used two different algorithms for clustering cells according to their transcriptomes of microinjected cells. In Seurat-based clustering, progenitors appeared as one cluster in the microinjection data set (**Fig. 4B**), which could be separated in APs and BPs using RaceID clustering in this data set (**Fig. 4C**). In the electroporated cells dataset, Seurat-based clustering resolved APs and BPs (**Fig. 4F**). We plotted several known marker genes for detailed cluster annotations (for control and EPZ, for both electroporated (**Supplementary Fig. 4A, B**) and microinjected cells (according to RaceID clustering) (**Supplementary Fig. 4C, D**). This allowed us to clearly identify APs (expressing *Hes5, Fabp7*, or *Sox2*), BPs (positive for *Neurog2, Eomes,* or *Insm1*), different clusters of neurons (*Neurod6, Dcx, Dpysl3,* or *Tubb3*), and endothelial cells (exclusively in the electroporated samples) (**Fig. 4B, C, F**).

Using the Seurat-based cell clusters, we plotted the cells for each of the two conditions (control and EPZ) and showed that both conditions contributed to each cluster (**Fig. 4D, G**). We concluded that all expected progenitor types were captured with both labelling approaches and in both conditions.

Interestingly, besides the main cell populations (APs, BPs, neurons) we identified one cell cluster lacking a known, specific transcriptional fingerprint (**Supplementary Fig. 4A, C**). We hypothesised that our high-resolution and clonal scRNA-seq approach captured a rare transient transcriptional state that we termed accordingly TTS (**Fig. 4B-H**). Few genes with enriched, but not unique, transcription were found in the TTS, to which belonged for example *Ofd1* and *Mme* (microinjection dataset) as well as *Fgfr3* and *Nr2f1* (electroporation dataset) (**Supplementary Fig. 4**). GO enrichment analysis (electroporation dataset) showed that genes up-regulated in the TTS compared to APs were enriched in terms associated with adhesion, lumen-formation and actin cytoskeleton elements (transcriptional signatures related to epithelial polarity; **Supplementary Fig. 5A,** left panel and **Supplementary Fig. 5B**). Genes with decreased expression in the TTS cluster compared to APs were enriched in GO terms associated with spindle and centrosome organisation, midbody, kinetochore and chromatin condensation (transcriptional signatures related to cell division; **Supplementary Fig. 5A**, right panel and **Supplementary Fig. 5B**). This transcriptional signature of TTS suggests that this cell state relates to APs (**Supplementary Fig. 5**), but that it has reduced proliferation capacity.

While the TTS comprised a minor fraction in control conditions, its proportion increased upon EPZ treatment, together with a decrease in the proportion of APs (**Fig. 4E, H**). The TTS’s transcriptional makeup described above had hallmarks of decreased proliferative capacity. Upon DOT1L inhibition we observed, e.g., an increased expression of *Fgfr3* in the TTS (**Supplementary Fig. 4B**). Increased levels of *Fgfr3* were reported to induce cell cycle exit of progenitors (Huang et al., 2020). Thus, the scRNA-seq data reflect at the transcriptional level our data on single-cell reconstruction of lineage progression, and both experimental attempts show a general increase in the neurogenic potential of APs upon DOT1L inhibition.

### DOT1L inhibition favours neuronal lineage progression by PRC2-derepression of metabolic genes

To elucidate the molecular mechanisms underlying changes in neuronal lineage progression and the switch of APs towards neurogenic division upon DOT1L inhibition, we performed unbiased analysis of (i) alterations of TF activities in gene regulatory networks, (ii) differentially expressed genes (DEGs) in APs and in the TTS, and (iii) GO terms enriched within DEGs, of electroporated cell scRNA-seq data. The analysis of changes in the TF network activity in response to DOT1L inhibition, in a cell-type resolved manner, revealed a set of key transcriptional regulators with cell-type specific activity changes (**Fig. 5A**). *Atf4, Cepbg, Rad21, Ezh2, Sox2, 4, 9, 21, Lhx2, Pou3f2,* and *Pou2f2* changed activity upon DOT1L inhibition in APs, TTS or BPs. Intriguingly, analysis of GO terms that enriched in APs and TTS, revealed that the DEGs upon DOT1L inhibition strongly related to metabolism and to oxidative phosphorylation (OxPhos), a hallmark of neuronal differentiation (**Fig. 5B**).

**Figure 5.**
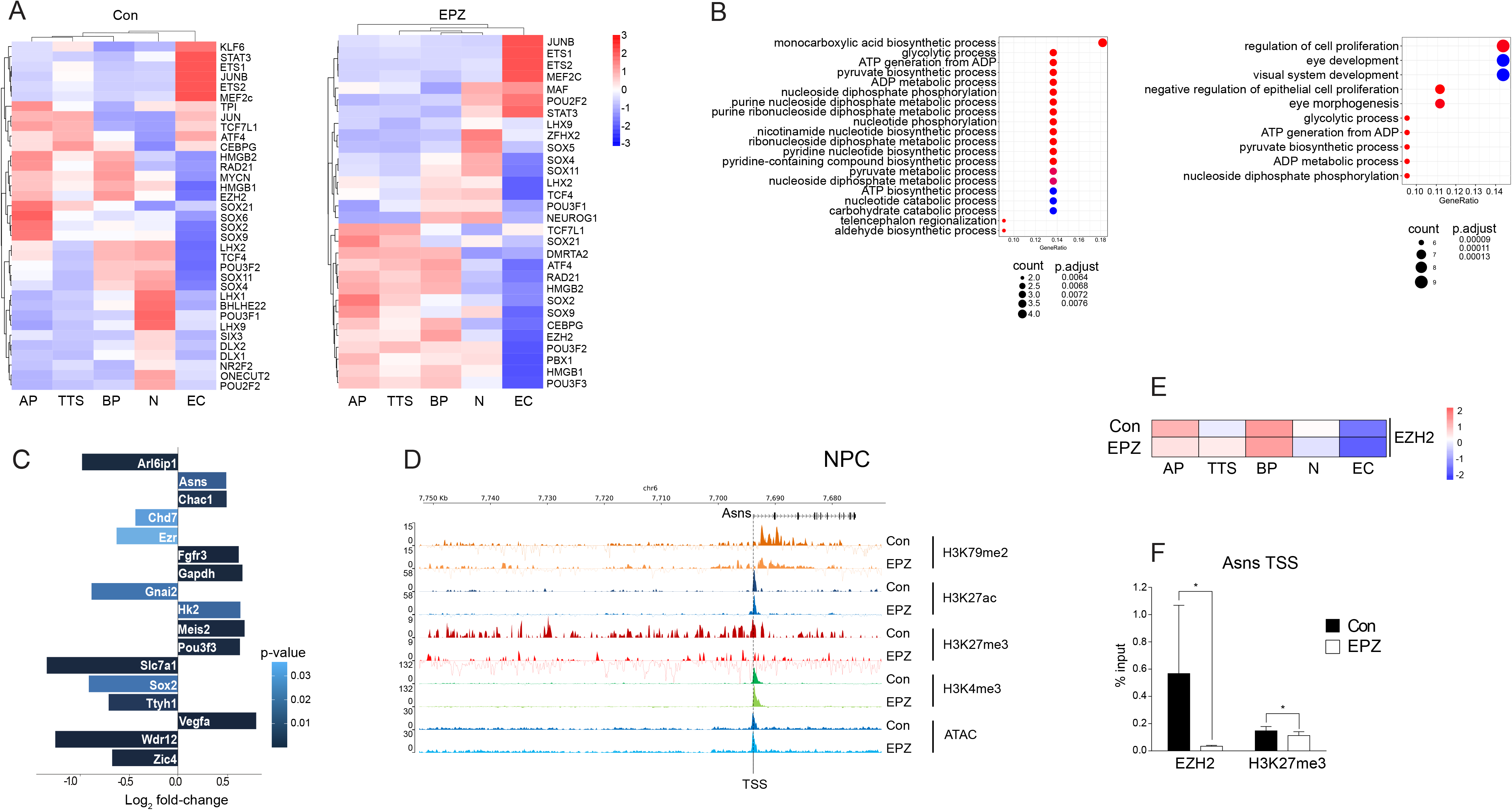
DOT1L inhibition leads to PRC2 de-repression of *Asns* gene locus in neural progenitors. **A)** Heatmap of results from SCENIC analysis showing activities of regulons and transcription factors controlling them in control (left) and EPZ (right) treated condition, resolved according to cell type/state. Scale represents enrichment scores for regulons expressed as area under recovery curve (AUC) of gene expression-value-based rankings across all genes for each cell. **B)** Left: Top GO terms (p.adjust < 0.00170) retrieved with genes increased in expression in TTS only (EPZ vs Con). Right: Top GO terms (p.adjust < 2e-09) retrieved with genes decreased in expression in APs only (EPZ vs Con). **C)** DEGs upon DOT1L inhibition in APs and TTS directly targeted by transcription factors captured in the SCENIC analysis. Given is the log_2_FoldChange of the respective DEG and as shades of blue the p.adjust value. **D)** ChIP-seq and ATAC-seq tracks showing distribution of peaks for selected chromatin marks on the *Asns* locus for NPCs. Tracks for control and EPZ conditions are compared. Indicated is the TSS of *Asns*. **E)** Comparison of the activity patterns of regulons controlled by EZH2 upon EPZ inhibition and in controls, resolved according to cell type/state. Scale as in A). **F)** ChIP-qPCR analyses for EZH2 and H3K27me3 at the TSS on *Asns* genomic locus in control and EPZ treated condition. One-sided paired Wilcoxon statistical test was performed, error bars represent SEM. *p<0.04.

We next merged the preceding analyses to identify potential DEGs that had predicted binding sites for the TF subset with altered activities upon DOT1L inhibition. This analysis retrieved 17 potential target genes, among which we found increased expression of three metabolic enzymes that are involved in producing intermediates that fuel, directly or indirectly, the TCA cycle, i.e. *Asns, Gapdh,* and *Hk2* (**Fig. 5C**).

In this set of target genes, the increased expression of *Asns* was our top candidate as misexpression of *Asns* is not only involved in leukaemia (Lomelino et al., 2017) (for which deregulated functions of DOT1L have been described (Okada et al., 2005)), but its mutation also causes microcephaly (Ruzzo et al., 2013). However, DOT1L is generally considered to favour transcription, thus increased expression upon its inhibition is seemingly counter-intuitive. To gain insight into the mechanism(s) that increased expression levels of *Asns* upon DOT1L inhibition, we made use of an extensive quantitative epigenetic dataset of mouse embryonic stem cells derived NPCs (Ferrari et al., 2020). The ChIP-seq profiles from NPCs at the *Asns* gene locus revealed specifically decreased levels of H3K27me3 alongside with decreased levels of H3K79me2 upon DOT1L inhibition (**Fig. 5D**). This epigenetic profile suggested that PRC2 de-repression resulted in increased expression levels of *Asns* upon DOT1L inhibition in NPCs. In line with this hypothesis, the gene regulatory network with altered activity upon DOT1L inhibition included EZH2 (**Fig. 5A, E**) in our tissue paradigm. ChIP-qPCR in NPCs confirmed that DOT1L inhibition reduced the level of H3K27me3 by reducing the presence of EZH2 at the TSS of the *Asns* gene locus (**Fig. 5F**). These data suggest the presence of a local regulatory network in which a lower activity of DOT1L results in lower activity of EZH2 at the *Asns* gene locus that causes less H3K27me3 at the TSS of *Asns,* increasing its expression. Our data thus provide a mechanistic link between DOT1L inhibition and increased expression of target genes.

### DOT1L affects lineage progression by regulating AP metabolism in an ASNS-dependent manner

To assess if *Asns* was functionally involved in the cellular phenotype we observed upon DOT1L inhibition, we examined if ASNS inhibition would rescue the EPZ-induced premature differentiation of APs. We labelled APs by *ex utero* electroporation of a mCherry reporter plasmid and incubated the hemispheres for 24 h with DMSO (control), or with EPZ either with or without L-Albizziine (L-Alb), an inhibitor of ASNS (**Fig. 6**). We assessed the extent of neurogenesis and differentiation using co-immunostainings for TUBB3 (**Fig. 6A**). In line with our previous data (**Fig. 2C, G**), we observed an increase in TUBB3-expressing cells in the electroporated cells upon DOT1L inhibition (**Fig. 6B**). The increased neuronal differentiation upon EPZ treatment was completely abolished by co-inhibition of the ASNS activity (**Fig. 6B**). We next examined if the altered expression of *Asns* might be a key driver of AP’s neuronal differentiation *per se*. We hence overexpressed ASNS together with a GFP reporter using *ex utero* electroporation and analyzed the TUBB3 expression in the electroporated cells (**Fig. 6C**). Upon overexpression of ASNS in APs, we observed twice as many TUBB3 positive cells compared to control (**Fig. 6D**). Thus, taken together, our data show that the differentiative role of DOT1L in APs is mediated by ASNS-dependent metabolic regulation.

**Figure 6.**
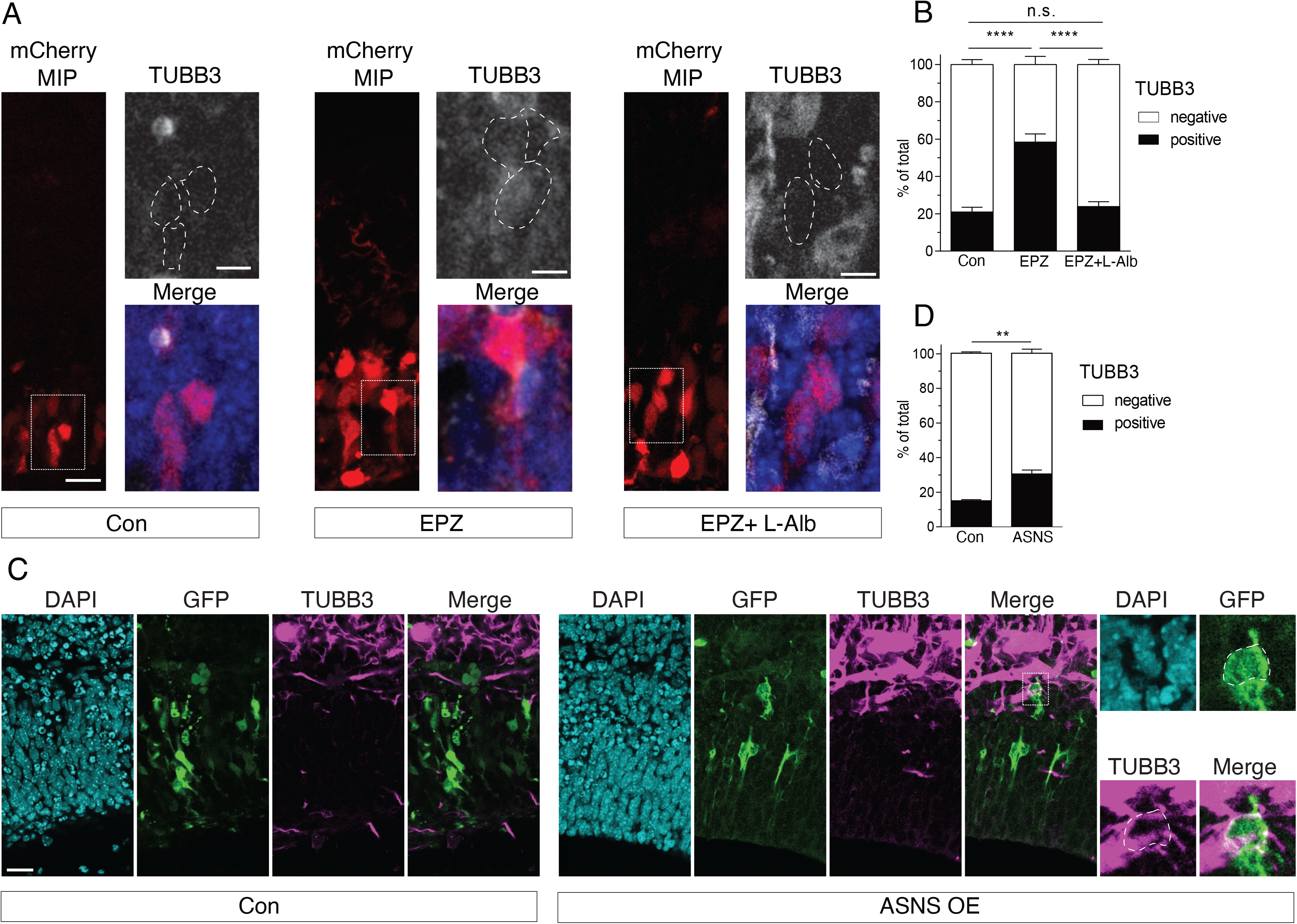
Increased ASNS expression mediates differentiative effects in response to DOT1L inhibition. **A)** Overview of mCherry+ cells in IF staining for TUBB3 in control (Con), EPZ, and co-inhibition (EPZ + L-Albizziine (L-Alb)) conditions. Scale bars in main panels 200 µm, in insets 50 µm. **B)** Fraction of TUBB3+ and TUBB3- cells expressed as percentage of total cells per condition. Quantifications were made from independent experiments (n=3), 183 cells in total (Con: 118 cells, EPZ: 182 cells, L-Alb/EPZ: 174 cells). Fisher’s exact test, ****p<0.0001, error bars represent SEM. MIP - maximum intensity projections. **C)** Electroporated cells expressing GFP (green) or TUBB3 (magenta) in control (Con; pCMV-GFP) or in ASNS overexpression (ASNS OE; pCMV-ASNS-GFP) condition 24 h after slice culture. Scale bars: 20 µm. **D)** Fraction of TUBB3+ and TUBB3- cells expressed as percentage of total cells per condition. Quantifications were made from independent experiments (n=3). Student’s t-test, **p<0.01, error bars represent SEM.

## Discussion

In this study, we investigate the impact of DOT1L enzymatic activity in regulating neural progenitor behaviour during neurogenesis by combining pharmacological inhibition of DOT1L with single-cell tracing in tissue, high-resolution lineage reconstruction and transcriptomic profiling of lineage-related cells. Our data suggest that DOT1L inhibition changes the mode of division of APs, the founder progenitor population of the neocortex, resulting in an increase in neuronal differentiation. Highly resolved lineage tracing and analysis shows that DOT1L inhibition favours symmetric consumptive division (AP➔N+N) over asymmetric self-renewing division (AP➔AP+BP). To the best of our knowledge, this is the first transcriptional regulator described altering AP’s division mode in this manner. Transcriptional profiling reveals that DOT1L influences fate transition and suggests it affects the symmetry/asymmetry of cell fate by regulating crucial metabolic enzymes. Our findings link the epigenetic landscape of neural progenitors, their metabolic state and fine cell biological processes leading to delamination and fate transition.

### DOT1L and the AP division mode

In our work, we make use of highly resolved cell biological analyses in tissue, and we focused our attention on the division mode, cellular output and delamination as they represent the different faces of a much broader repertoire of processes defining cell identity and driving fate transition. Extending on previous findings (Franz et al., 2019; Ferrari et al., 2020), our present data strongly suggest that DOT1L activity preserves asymmetric self-renewing divisions of APs and in turn prevents delamination. How DOT1L affects the asymmetry of division and cell delamination remains still an open question. One possibility might be that DOT1L favours the generation of an asymmetric progeny (one AP and one BP) by placing asymmetric H3K79 methylation marks at the chromatin of one of the two daughter cells. Despite epigenetic features being recognised as an inheritable information affecting cell fate, genome-wide data of asymmetric epigenetic inheritance is rare and difficult to achieve in mammalian cells (Xie et al., 2017; Ma et al., 2020). Our data present an indication that H3K79 methylation might serve as epigenetic memory of an AP fate and that DOT1L activity is needed to keep this progenitor state. Given that H3K79 methylation is a relative stable histone mark, erased rather through cell division than demethylation (Chory et al., 2019), a memory function of H3K79 methylation would be an ideal feature.

### DOT1L/PRC2 epigenetic crosstalk regulates the choice between self-renewal and differentiation in APs

Mechanistically, the study presented here provides insight into a local DOT1L/PRC2 crosstalk reminiscent of the local epigenetic signature associated with promoters of genes with increased expression upon DOT1L inhibition in NPCs *in vitro* (Ferrari et al., 2020). In NPCs, genes that transcriptionally increase upon DOT1L inhibition associate with bivalent or PRC2 repressed promoters. Of note, the use of EPZ uncouples DOT1L activity from its scaffolding functions and therefore allows us to link the molecular and cellular phenotype(s) with the methyltransferase activity of DOT1L. We present here evidence that DOT1L enzymatic activity affects PRC2 activity in cortical tissue. DOT1L inhibition leads to reduced activity of the PRC2 member EZH2 in APs compared to control.

We propose that, in control condition, higher EZH2/PRC2 activity in APs and BPs compared to neurons is associated with maintenance or re-establishment of progenitor identity; in turn lower activity levels lead to neuronal differentiation. In line with this hypothesis, APs with *Eed* or *Ezh2* loss-of-function were found to have an accelerated and precocious cell cycle exit (Pereira et al., 2010; Telley et al., 2019).

Our data contribute to strengthen the concept that PRC2 activity is of major importance for preserving AP identity and self-renewal potential. Of note, regarding the crosstalk with PRC2, DOT1L inhibition reduced binding of EZH2 and H3K27me3 levels in NPCs *in vitro*.

Interestingly, DOT1L/PRC2 crosstalk takes place in other cell types, in which transcripts increase upon DOT1L inhibition (Aslam et al., 2021). In immune cells the proposed underlying mechanism is transcriptional decrease of *Ezh2* upon DOT1L loss-of-function (Aslam et al., 2021). In our lineage resolved scRNA-seq data, however, *Ezh2* transcription did not change upon DOT1L inhibition in APs, TTS, or BPs (data not shown), but its activity was altered. DOT1L/PRC2 crosstalk might therefore have different facets depending on the cell type.

### DOT1L links the epigenetic landscape to metabolism *via* ASNS

Metabolism has emerged as a crucial regulator of stem and progenitor cells. We here describe a previously unknown link between metabolism, epigenetics and APs by identifying *Asns* as a DOT1L/PRC2 target and crucial partner in the regulative network affecting AP fate choice and lineage progression.

*Asns* encodes for asparagine synthetase, an enzyme that converts glutamine (Gln) and aspartate (Asp) to glutamate (Glu) and asparagine (Asn). Glu can be further metabolised to 2-oxo-glutarate, feeding the TCA cycle and fostering mitochondrial energy supply through OxPhos (Zheng et al., 2016; Shiratori et al., 2019). ASNS thus balances not only the abundance of important amino acids used for protein synthesis but also feeds into other metabolic processes including the TCA cycle. The importance of the TCA cycle and mitochondrial metabolism is highlighted by recent findings showing that ARHGAP11B, a human specific gene involved in cortical expansion, mediates glutaminolysis and increases the Glu levels that feed the TCA cycle (Namba et al., 2020).

Intriguingly, ASNS dysfunction is linked to microcephaly (Ruzzo et al., 2013; Schleinitz et al., 2018). Of note, *Asns* loss-of-function (Ruzzo et al., 2013) as well as *Asns* overexpression (this study) seems to produce microcephalic features. *Asns* expression is lower in progenitors compared to neurons, both in human (Johnson et al., 2016) and in mice (Ruzzo et al., 2013). Thus, loss-of-function might affect neurons much more than progenitors, while overexpression might affect primarily APs, where low levels of *Asns* might be necessary to maintain their self-renewing potential.

The precise mechanisms linking DOT1L and ASNS activity to the regulation of AP behaviour are not yet clear. Based on the current literature, several options can be envisaged. DOT1L inhibition might alter Glu and/or Asp levels. Both amino acids can be metabolised to intermediates of the TCA cycle (2-oxo-glutarate, oxaloacetate), the activation of which could result in mitochondrial energy supply (OxPhos). OxPhos is the main energy source of cancer cells (Shiratori et al., 2019), and also of neurons (Zheng et al., 2016). In addition to its role as a metabolite, Glu was reported to regulate stem and progenitor cells when released in the extracellular space, where it depolarizes stem and progenitor cells, inhibiting DNA synthesis and favouring neurogenesis (LoTurco et al., 1995). Of note, Asn could also have a role in itself, by activating the mTOR pathway and/or acting as a sensor of specific cellular states or cell growth as reported for cancer cells (Krall et al., 2016; Krall et al., 2021).

Taken together, our study shows that DOT1L is a key regulator of cortical neurogenesis by preserving the self-renewing potential of APs. Mechanistically, DOT1L provides a link between the epigenetic landscape and Asn/Glu cell metabolism. The highly resolved lineage tracing showing the premature switch to symmetric consumptive divisions also provides a likely cell biological explanation for microcephalic phenotypes associated with *ASNS* (Ruzzo et al., 2013) and *DOT1L* mutations.

Placing our work in a broader frame, it is interesting to note that while primary microcephaly mutations in *ASPM* or *MCPH1* are associated with premature switch from symmetric proliferative to asymmetric self-renewing division (Fish et al., 2006; Gruber et al., 2011), DOT1L inhibition is the first one reported to be associated with the premature switch from asymmetric self-renewing to symmetric consumptive division. This premature switch is likely to decrease the number of BPs generated, further reducing the total neuronal output. These observations highlight the role of symmetry vs asymmetry of AP’s fate choice in defining the window of cortical neurogenesis and in affecting cortical development in physiology and pathology.

## Material and Methods

### Mice

All animal studies were conducted in accordance with German animal welfare legislation, and the necessary licenses were obtained from the regional Ethical Commission for Animal Experimentation of Dresden, Germany (Tierversuchskommission, Landesdirektion Dresden) or the Regierungspräsidium Freiburg (X-17/03S). Wild type mice (C57BL/6J) were harvested at E14.5. Where specified, microinjection experiments were performed with *Tis21*-*Gfp* knock-in E14.5 mouse embryos. To obtain *Tis21*-*Gfp* knock-in mice, homozygous *Tis21*-*Gfp* knock-in male mice, Tis21+/tm2(Gfp)Wbh (Haubensak et al., 2004), were mated with C57BL/6J females.

### Dissection and organotypic slice preparation

Mice pregnant with E14.5 embryos were sacrificed via cervical dislocation for litter harvesting. Embryos were transferred into ice-cold phosphate buffered saline (PBS), decapitated, and heads collected in ice-cold PBS for *ex utero* electroporation. For microinjection, heads of embryos were dissected in pre-warmed Tyrode’s solution (37°C) for organotypic slice preparation as previously described (Taverna et al., 2012; Wong et al., 2014). 250 µm vibratome sections were prepared and transferred into pre-warmed slice culture medium (SCM) (Taverna et al., 2012). The composition of the SCM is as follows: Neurobasal medium (Thermo Fisher Scientific, Germany), 10% rat serum (Charles River, Japan), 2 mM L-glutamine (Thermo Fisher Scientific), Penstrep (Thermo Fisher Scientific), N2 supplement (Thermo Fisher Scientific), B27 supplement (Thermo Fisher Scientific), 10 mM Hepes-NaOH pH 7.3. Before the start of microinjection, brain slices were transferred to 3.5 cm dishes containing 37°C warm CO_2_-independent microinjection medium (CIMM: DMEM-F12 (Sigma, Germany, D2906), 2 mM L-glutamine, Penstrep, N2 and B27 supplements, 25 mM (final concentration), Hepes-NaOH pH 7.3).

### Microinjection and slice culture

Microinjection into single APs in tissue was performed on organotypic slices from E14.5 developing mouse telencephalon. Depending on the purpose of the experiment, we used either manual or automated microinjection. Manual microinjection into single APs in tissue was used for lineage tracing experiments, following previously described protocols (Taverna et al., 2012; Wong et al., 2014). Automated microinjection into single APs in tissue was used for scRNA-seq experiments, where a higher throughput was needed. Automated microinjection was performed using the Autoinjector 1.0, a recently developed high-throughput robotic platform (Shull et al., 2019; Shull et al., 2021). For all experiments, microinjection was performed in pre-warmed CO_2_-independent microinjection medium (CIMM) (Taverna et al., 2012). Briefly, 1.5 µl of injection solution (5 µg/µl Dextran-Alexa488 or Dextran-Alexa555) was loaded into a microcapillary needle for microinjection. The microinjection needle was mounted on the capillary holder and microinjection was performed, either manually or with Autoinjector 1.0. by approaching the ventricular (apical) surface of organotypic tissue slices. Microinjected slices were embedded in collagen matrix that was allowed to solidify at 37°C for 5 min. After this time, each dish received 2 ml of either SCM containing 0.09% dimethyl sulfoxide (DMSO, Control, Con) or 9 nM DOT1L inhibitor Pinometostat (EPZ5676, EPZ) (Selleckchem, USA) and slices were transferred and kept in culture at 37°C in a humidified atmosphere of 40% O_2_ / 5% CO_2_ / 55% N_2_. Unless stated otherwise, organotypic slices were kept in culture for 24 h, a time window corresponding to one complete cell cycle of APs at mid-neurogenesis (Arai et al., 2011).

### *Ex utero* electroporation and hemisphere rotation culture

*Ex utero* electroporation of E14.5 mouse embryos was performed in sterile PBS. An intraventricular injection of a solution containing 1-1.5 µg/µl pCAG-GFP, pCAG-mCherry, pCMV-GFP (PS100010, Origene, USA) or pCMV-ASNS-GFP (MG208883, Origene) mixed with 0.1% Fast green in sterile PBS was performed. It was immediately followed by 5-6 pulses of 28-30 V, 50 ms each at 1 s intervals delivered through platinum tweezer electrodes (3 mm diameter) using a BTX ECM830 electroporator, similarly to what previously described (Calegari et al., 2002). Subsequently, electroporated hemispheres were dissected and placed in flasks containing 2 ml of SCM, where indicated with either EPZ or DMSO, for 24 h at 37°C and gentle rotation in a humidified atmosphere of 40% O_2_ / 5% CO_2_ / 55% N_2_. At the end of the culture, hemispheres were washed in 1X sterile PBS to remove media and processed for immunohistochemistry, or dissociated and processed by FACS followed by scRNA-seq. For ASNS/DOT1L co-inhibition experiments, electroporation of pCAG-mCherry was performed as described above using a NEPA21 super electroporator (NEPAGENE, Japan). Electroporated hemispheres were dissected and placed in glass culture dishes containing 2 ml of SCM with three conditions: EPZ only, EPZ in combination with 4 mM L-Albizziine (GoldBio, USA), or DMSO only. At the end of the culture, hemispheres were washed in 1X sterile PBS, fixed and processed for immunofluorescence analysis.

### Tissue dissociation and FACS sorting

Following hemisphere rotation culture, the cortices were dissected. The electroporated area was identified and isolated using an epifluorescence microscope, and dissociated into single-cell suspensions using Neural Tissue Dissociation kit (P) (Miltenyi Biotec, Germany), following the manufacturer’s instructions. Microinjected slices were dissected, the microinjected area was identified and isolated using an epifluorescence microscope, then dissociated for sorting with the same dissociation kit as described above. Enrichment of labelled cells was performed via FACS on a BD FACS Diva 8.0.2 (**Supplementary Fig. 3B**). Cells were sorted into 384-well plates containing 240 nl lysis buffer (Herman et al., 2018). Sorted plates were centrifuged at 2000 X g for 1 min at 4°C and immediately transferred into a −80°C freezer until processing for scRNA-seq.

### Library preparation, single-cell RNA sequencing and data accessibility

Processing of sorted cells for scRNA-seq was performed as described (Hashimshony et al., 2016) with modifications based on the mCEL-Seq2 protocol (Herman et al., 2018). Briefly, 384-well plates containing sorted cells in lysis buffer were incubated for 3 min at 90°C and quickly cooled down to 4°C. cDNA was synthesized from RNA in each well by adding 160 nl of reverse transcription mix (Herman et al., 2018), followed by incubation for 1 h at 42°C. Heat inactivation of the reaction mix was performed for 10 min at 70°C. Synthesis of cDNA second-strand was performed at 16°C for 2 h. cDNA from 96 wells was pooled for sample clean up and *in vitro* transcription. This resulted in four sequencing libraries from a single 384-well plate. Libraries were sequenced (paired-end) on Illumina Hi-seq 2500 at a depth of ∼150,000 reads per cell. Raw scRNA-seq data along with expression matrices have been submitted to the online repository Gene Expression Omnibus (GEO), accessible via GSE176323.

### Fixation, embedding, and sectioning

Fixation, embedding, and sectioning were performed as described (Taverna et al., 2012; Gray de Cristoforis et al., 2020). Hemispheres or slices were washed twice with PBS to remove culture media, followed by fixation with 4% paraformaldehyde in 120 mM sodium phosphate buffer pH 7.4 at room temperature for 30 min followed by 4°C overnight. Following fixation, hemispheres and slices were washed thrice with 1X PBS. Coronal sections (50 µm) were prepared from microinjected slices on a vibratome (Leica VT1000S). All slices were collected in PBS and stored at 4°C for subsequent immunofluorescence staining. Electroporated hemispheres were infiltrated with 30% sucrose at 4°C overnight prior to embedding in TissueTek tissue freezing medium (Leica Biosystems, Germany). 16 µm-thick sections were cut using a cryostat (Leica Biosystems) and resulting sections were stored at −20°C for further processing.

### Immunofluorescence staining

Floating sections were processed for immunofluorescence (IF) staining as previously described (Taverna et al., 2012). In brief, sections were permeabilized with 0.3% Triton-X100 (Carl Roth, Germany) in PBS for 30 min, followed by quenching in 0.2 M glycine buffer pH 7.4 for 30 min. The sections were washed three times with IF buffer, and incubated with primary antibodies diluted in IF buffer overnight at 4°C. Subsequently, sections were washed 5 times for 5 min each time with IF buffer. Secondary antibodies were diluted in IF buffer with DAPI (Carl Roth, Germany) as nuclear counterstain. Sections were incubated at room temperature for 1 h followed by washing (5 times 5 min each) with IF buffer and then PBS. Stained sections were mounted in Mowiol (Sigma, Germany) and imaged. Cryosections were processed for IF staining as follows. Sections were dried for 5 min in a chemical hood then washed twice with 1X PBS, followed by permeabilization with 0.3% Triton-X100 (Carl Roth, Germany) in PBS for 30 min. Subsequently, sections were incubated with blocking solution (10% Normal Donkey Serum, in 0.3% TritonX-100/PBS) for 1 h at room temperature in a humidified chamber. Primary antibodies diluted in blocking solution were incubated with the sections overnight at 4°C, followed by washing 5 times for 5 min each time with 0.3% Triton-X100 in PBS. Sections were then incubated with secondary antibodies diluted in blocking solution with DAPI as nuclear counterstain for 1 h at room temperature followed by washing for 5 times 5 min each time.

A complete list of all primary and secondary antibodies used together with corresponding concentrations are listed in table 1.

### Microscopy and figure preparation

Imaging was performed with confocal (Leica TCS SP8, Leica, Germany; Zeiss LSM 780 NLO; Zeiss, Germany) or widefield fluorescence (Zeiss Axio Imager M2; Zeiss) microscopes. Images shown are 1 µm-thick single optical sections, unless stated otherwise. Confocal images were taken at either 63X or 40X magnifications. Immunofluorescence images of single optical sections were quantified using Fiji (Schindelin et al., 2012). Final image panels were processed with Inkscape 1.0.2, Adobe Illustrator CS5.1 or Affinity publisher 1.10.5.

### Quantification procedure for *ex vivo* lineage tracing

We used a combined panel of cell morphological parameters and marker expression to score labelled cells and progeny in tissue.

Microinjected cells and their progeny were identified as cells positive for the microinjection dye upon inspection at the epifluorescence microscope. To assess cellular and morphological organization and marker expression, all positive cells in an experiment were imaged at high-resolution using confocal microscopy (see also above). Microinjected cells and their progeny were scored for presence or absence of apical contact by matching the signal of injected cells and their morphology with the general tissue structure, revealed by DAPI staining (presence of contact: apical contact+; absence of contact: apical contact-). Symmetry (apical contact+/apical contact+; apical contact-/apical contact-) versus asymmetry (apical contact+/apical contact-) of apical contact in daughter cell pairs was assessed. Numbers of cells with or without apical contact were calculated as a percentage of total microinjected cells scored.

Positive microinjected cells were scored for the immunoreactivity to the cytoskeletal protein tubulin beta 3 class III (TUBB3) and classified as negative (TUBB3-) or positive (TUBB3+). Proportions of TUBB3+ and TUBB3- were calculated as percentage of total microinjected (or electroporated for the electroporation experiments) cells per condition. The BP marker Eomesodermin (EOMES) was used to score BP fractions arising after culture. Cells positive for EOMES were quantified as EOMES+ and those negative for it marked as EOMES-. Proportions of EOMES+ and EOMES- cells were calculated as for TUBB3, see above.

To assess symmetry versus asymmetry of cell lineages arising from APs, daughter cell pairs were assessed for expression of TUBB3 and EOMES as described above. Daughter cell pairs were scored as symmetric for TUBB3 expression, meaning both cells were positive for TUBB3 (TUBB3+/TUBB3+) or negative (TUBB3-/TUBB3-), or asymmetric for TUBB3 expression (TUBB3+/TUBB3-). Proportions of each scored group were calculated as a percentage of total daughter cell number. The same approach was used to score symmetric versus asymmetric inheritance of BP identity (symmetric: EOMES+/EOMES+, EOMES-/EOMES-; asymmetric: EOMES+/EOMES-). All lineage tracing experiments were performed on a minimum of three biological replicates.

### ChIP-qPCR from neural progenitor cells

ChIP-qPCR was performed from neural progenitor cells (NPCs) as previously described (Bovio et al., 2019; Ferrari et al., 2020), with minor changes in composition of buffers: Lysis buffer for EZH2 (10 mM EDTA; 50 mM TRIS-HCl pH 8; 0,2 % sodium dodecyl sulfate (SDS) and 1X protease inhibitor), lysis buffer for H3K27me3 (10 mM EDTA; 50 mM TRIS-HCl pH 8; 1% SDS and 1X protease inhibitor), dilution buffer (20 mM TRIS-HCl; 2 mM EDTA; 150 mM NaCl and 1% Triton). All ChIP-qPCR experiments were performed with NPCs, generated from V6.5 mESCs as described (Ferrari et al., 2020), cultured on 1 x 10 cm cell culture dishes and treated for 48 h with 10 µM EPZ or 0.1% DMSO as control. Approximately 6 million cells were used in each ChIP experiment. 5 µg of either H3K27me3 (#39155, Active Motif, USA) or 5 µg EZH2 (#39901, Active Motif) antibody was used. Significant differences in H3K27me3 and EZH2 binding to the selected locus were calculated between EPZ and control using one-sided paired Wilcoxon statistical test. Results from ChIP-qPCR experiments are from a minimum of five independent replicates. The following primers targeting the transcriptional start site (TSS) on *Asns* were used:

ASNS_TSS_fw: TCCCGCTTACCTGAGCACTA

ASNS_TSS_rv: CAGCCACATGATGAAACTTCC

### Bioinformatics analysis

All bioinformatics analysis are reproducible and stored as R scripts and markdown files on the online repository Github, accessible via: https://github.com/Vogel-lab/DOT1L_activity_neocortex-paper

### Alignment, transcript quantification, and single-cell gene expression matrix

Alignment of paired-end reads to the murine transcriptome (Encode VM9 release, downloaded from the UCSC genome browser) was done using bwa (v0.6.2-r126) as previously published (Li and Durbin, 2009) with default parameters. Different isoforms of the same gene were merged onto one gene locus. Reads mapping to multiple loci were excluded. A total of 750 cells (Con: 371, EPZ: 379) were recovered from *ex utero* electroporation experiments for sequencing. Cells from two microinjection experiments were pooled. The first experiment recovered 213 (Con: 59, EPZ: 154) cells. More slices were injected for the second experiment, which increased the number to 533 recovered cells (Con: 248, EPZ: 285). Cells from both microinjection experiments were pooled. A total of 746 (Con: 307, EPZ: 439) cells were recovered from the two microinjection experiments combined. In both *ex utero* electroporation and microinjection experiments, 34205 genes were retrieved with 7 and 18 cells not passing the sequencing, respectively, due to low read quality. Batch effects were negligible in the pooled cells from the microinjection experiments, as the cells from both experiments clustered together and were represented in all clusters. Transcripts were assigned to single-cells using information from barcodes on the left read, with the first six bases representing UMI and the next six bases corresponding to cell specific barcodes (Herman et al., 2018). Counts for each transcript per cell were obtained by aggregating the number of UMIs per transcript mapping to each gene locus, and based on binomial statistics, the number of observed UMIs was converted into transcript counts (Grün et al., 2014). Subsequently, a single-cell transcript count matrix was constructed with row and column names corresponding to genes and cells, respectively.

### scRNA-seq data analysis with Seurat v3.0

All analyses with Seurat v3.0 (Stuart et al., 2019) were done separately for microinjection and electroporation datasets in R (v3.6.1) and Rstudio (v1.2.5042). Count matrices for microinjection and electroporation datasets were loaded and converted into SingleCellExperiment objects. Additional Bioconductor packages were used in combination with Seurat. Data processing steps and parameters, unless stated otherwise, were kept the same for both datasets. All data pre-processing and clustering steps were performed on a combined matrix containing cells from both conditions (Con and EPZ). Cells from the different experimental technique (electroporation / microinjection) were analyzed separately but with Con and EPZ cells combined into one matrix. All genes with row sums greater than zero were kept as expressed genes. Subsequently, total counts and distribution of counts for cells in both control and EPZ conditions were calculated. Quality control was performed using Scater (McCarthy et al., 2017) resulting in 17456 and 17192 expressed genes for electroporation and microinjection datasets, respectively. Cells were filtered by excluding all those with a library size below 2500 total transcript counts. This excluded 15 cells (electroporation) and 89 cells (microinjection) from further analysis. The next filtering step excluded all cells with expressed features (genes) below 1000 (i.e. 1 cell from electroporation and 29 cells from microinjection). Cells with outlier values for mitochondrial genes were identified based on the median absolute deviation from the median number of mitochondrial genes across all cells. This filtering step discarded 16 cells (electroporation) and 21 cells (microinjection), retrieving 711 (Con: 350, EPZ: 361) and 589 (Con: 248, EPZ: 341) high quality cells for clustering analysis in electroporation and microinjection datasets, respectively.

800 of the highly variable genes were selected for computing principal components. The top eight principal components (PC) were used for clustering cells from electroporation and microinjection datasets, respectively. For the top eight PC both data sets showed strong enrichment of low p-value features as suggested by a drop of the p-value from the JackStraw plot and the visible elbow in the Elbow plot (**Supplementary Fig. 6A, B**). Unbiased clusters were annotated to cell types using a combination of top expressed features and published markers for cell types. In the microinjection dataset, AP and BP clusters were not resolved as separate clusters using Seurat. Thus, the resolution parameter was varied between 0.4 to 1.2 in increments of 0.1 to achieve optimized clustering. The resulting clusters were compared to each other using the adjusted rand index (ARI). From the average silhouette and the ARI, a resolution of 0.4 seemed to be most stable for both microinjection and electroporation datasets (**Supplementary Fig. 6A, B**). Increasing the number of PCs used for clustering to 10 in the microinjection dataset did not resolve AP/BP clusters (**Supplementary Fig. 6C**). Increasing the resolution to 1.2 in the microinjection dataset resulted in additional clusters created from the initial ones, but did not separate the AP/BP cluster (**Supplementary Fig. 6D**). Because of this limitation, microinjected scRNA-seq data were analyzed with RaceID3 (see below). Differential expression analysis was done based on standard settings in Seurat. Fisher’s exact test was computed on the contribution of treatment conditions to each cluster.

### scRNA-seq data analysis with RaceID3

scRNA-seq data from the microinjection experiment (728 cells) was analysed also with RaceID3, with minor adaptations to the default parameters (Herman et al., 2018). Before the filtering step, low quality cells as previously identified (Grün et al., 2016) were discarded. Filtering was done with the following parameters: mintotal = 2500, minexpr = 10, minnumber = 5, ccor = 0.4. Subsequent steps in RaceID3 analysis were run with the following parameters: metric = spearman, cln = 5, sat = False, outlg = 10. Marker genes for each cluster, intra-cluster differentially expressed genes (EPZ versus DMSO control), as well as global differentially expressed genes between EPZ versus DMSO control were extracted using default settings in RaceID3. Following quality control and filtering of scRNA-seq data from the microinjection experiment, 639 cells (DMSO: 264, EPZ: 375) passed the quality control and filtering step for clustering and further downstream analysis. Fisher’s exact test was computed on the contribution of treatment conditions to each cluster.

### Single-cell regulatory network inference and clustering (SCENIC) analysis

Gene regulatory network activity (GRN) in each cell was computed as previously described (Aibar et al., 2017). This tool takes Seurat objects as input. Seurat objects were exported after clustering analysis of the electroporation dataset and divided into separate objects for control and EPZ cells, respectively. Default settings were used for building the co-expression network and creating regulons. The top 50% regulons were used in the GRN activity scoring. Heatmaps of the GRN activities per cell were generated using pheatmap (v1.0.12). Since each GRN is controlled by one TF, GRN activity and TF activity is used interchangeably here. TFs with changed patterns of activity upon DOT1L inhibition were extracted along with their target genes, which they are predicted to bind to with high conformity. Subsequently, those target genes were intersected with DEGs from each cluster and this list was kept as TF target genes with altered expression upon DOT1L inhibition.

### Gene ontology (GO) term analysis

Over-representation analysis was performed using the enrichGO function of ClusterProfiler (v3.99.1). The following parameters were used: OrgDB = org.Mm.eg.db, ont = CC (for cellular components) or BP (for biological process), pAdjustMethod = BH, pvalueCutoff = 0.01, qvalueCutoff = 0.05, readable = TRUE. Subsequently, dot plots of the top GO terms were made ordering genes by gene ratio and adjusted p-values.

### ChIP- and ATAC-seq analysis

ChIP- and ATAC-seq analysis was performed on previously published data, gene expression omnibus (GEO) series accessions numbers GSE135318 for H3K79me2, H3K27me3, H3K4me3, and ATAC in NPCs (Ferrari et al., 2020) with published RPKM normalized reads. The mean of individual replicates was calculated with deepTools2 bigwigCompare with a bin size of 25 bp before subtracting the mean input signal from the corresponding mean ChIP signal. DeepTools2 pyGenomeTracks v3.5 was used to plot the reads over the displayed genomic regions (*Asns* chr6:7,670,648-7,751,776).

## Acknowledgement

The authors thank Francesco Ferrari for generating the used ChIP-seq data sets, and Annalisa Izzo for ChIP-qPCR expertise. We also thank the entire Vogel lab for discussions and critical comments on the manuscript. We thank Sagar for help with sequencing. We thank Mareike Albert and Takashi Namba for the critical comments on the manuscript and fruitful discussion on the data. We are grateful to the Services and Facilities of the Max Planck Institute of Molecular Cell Biology and Genetics for the outstanding support provided, notably Jussi Helppi and his team of the Animal Facility, Jan Peychl and his team of the Light Microscopy Facility, and Julia Jarrels and Ina Nusslein of the FACS facility.

This study was supported by the German Research Foundation 322977937/GRK2344 (BA, CF, TV, DG, MT, HB) and SPP1937/GR4980/1 (DG, PZ).

---

**Supplementary Figure 1.**
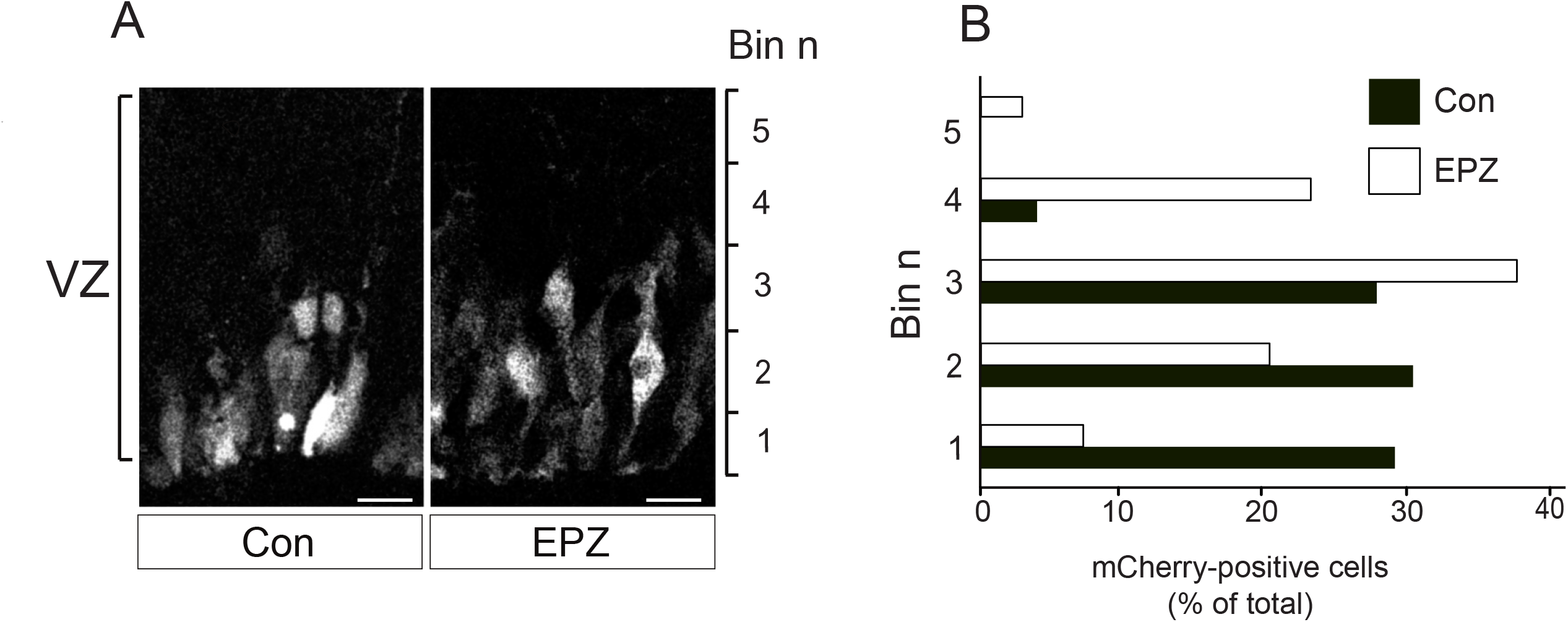
Basal distribution of mCherry-positive electroporated cells upon DOT1L-inhibition. **A)** Electroporated tissue section showing mCherry-positive cells (grey) in VZ of control (Con) and EPZ conditions after 24 h culture, binning of the VZ is indicated on the right side. **B)** Graph showing percentage of control and EPZ treated cells in binned distances from apical surface. Distance of Dx-A555 labelled cells from apical surface was divided into 5 bins of 30 µm intervals. Scale bar: 10 µm.

**Supplementary Figure 2.**
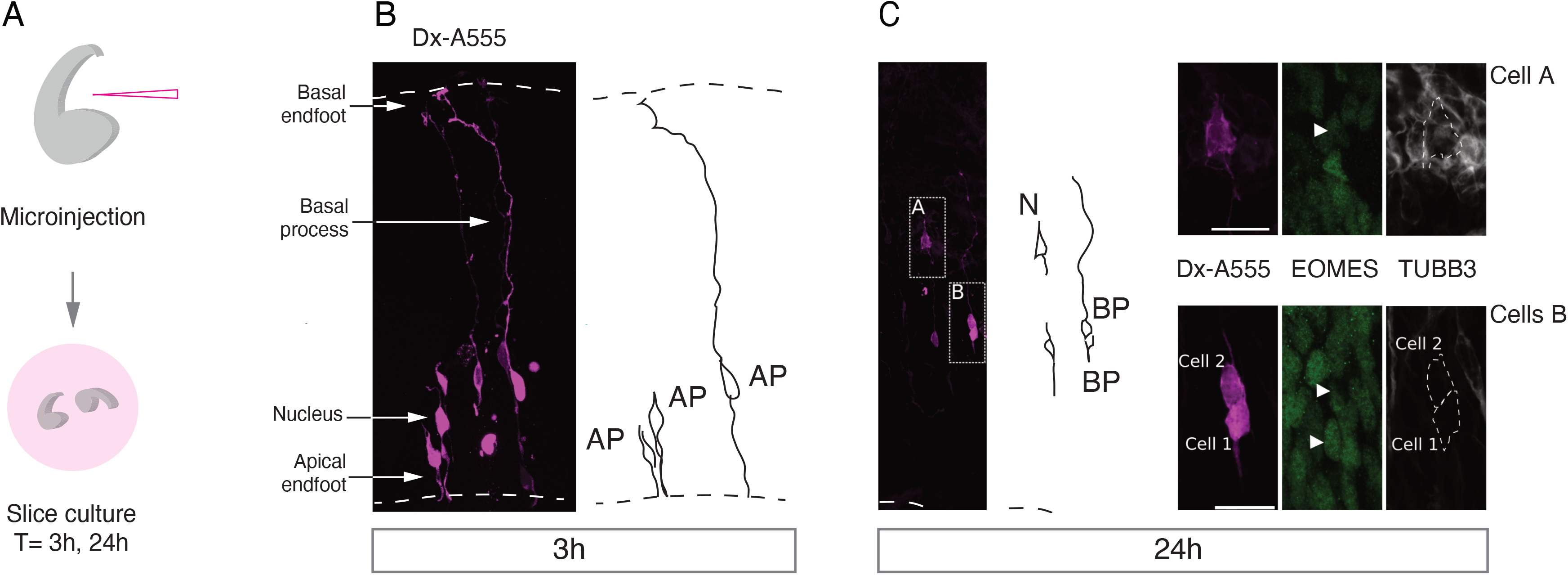
Additional information on microinjection and lineage tracing experimental procedure. **A)** Outline of microinjection and slice culture. **B)** Overview of microinjected tissue section showing Dx-A555 labelled cells after 3 h slice culture. **C)** Microinjected cells and progeny expressing EOMES and/or TUBB3 after 24 h slice culture.

**Supplementary Figure 3.**
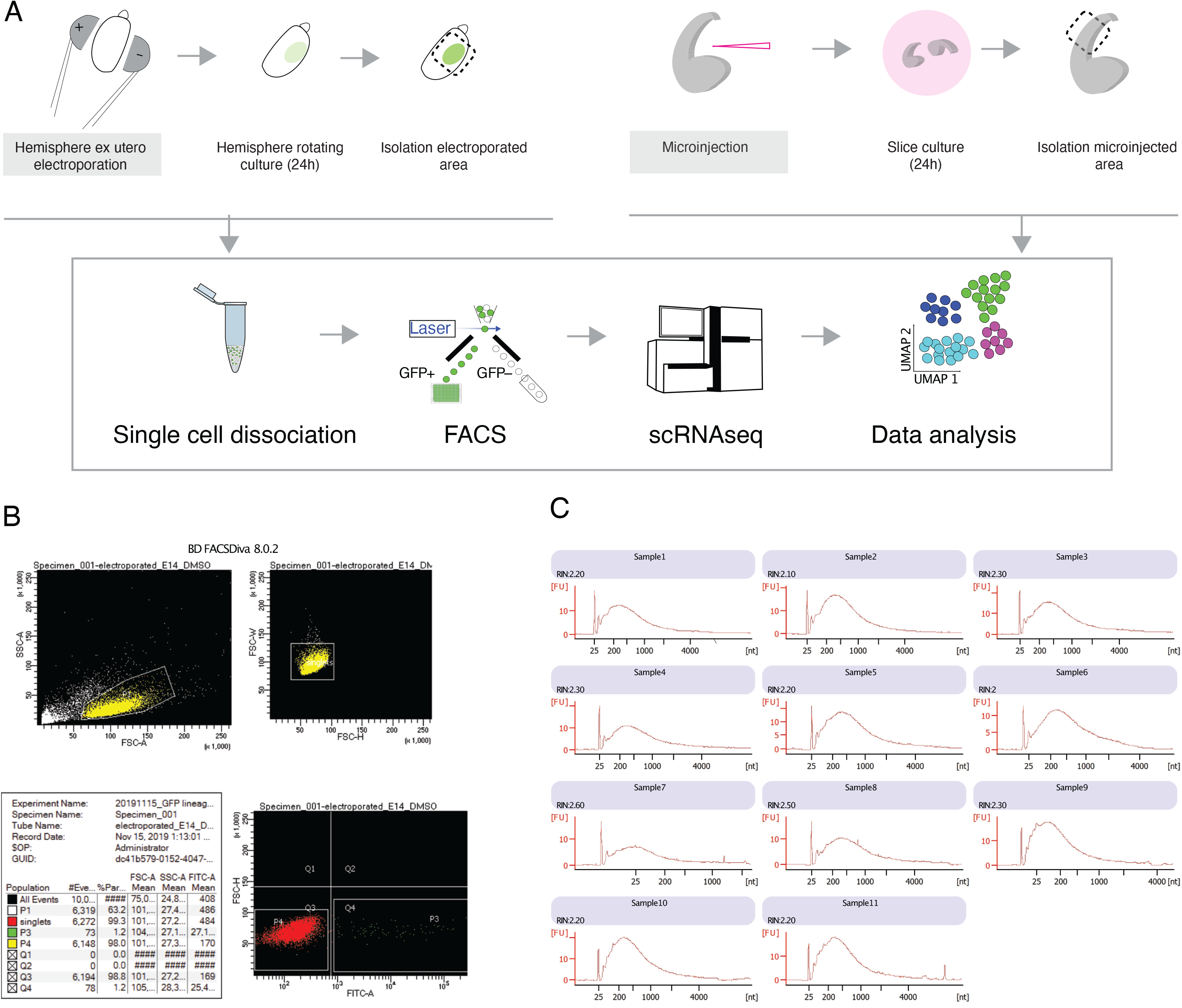
Additional information on sample acquisition for single cell RNA sequencing. **A)** Outline of procedure for scRNA-seq sample and data acquisition. **B)** Overview of gating and sorting parameters used for acquiring labelled cells. **C)** Snapshot showing concentration and fragment size distribution of RNA used for next generation sequencing (NGS) library preparation.

**Supplementary Figure 4.**
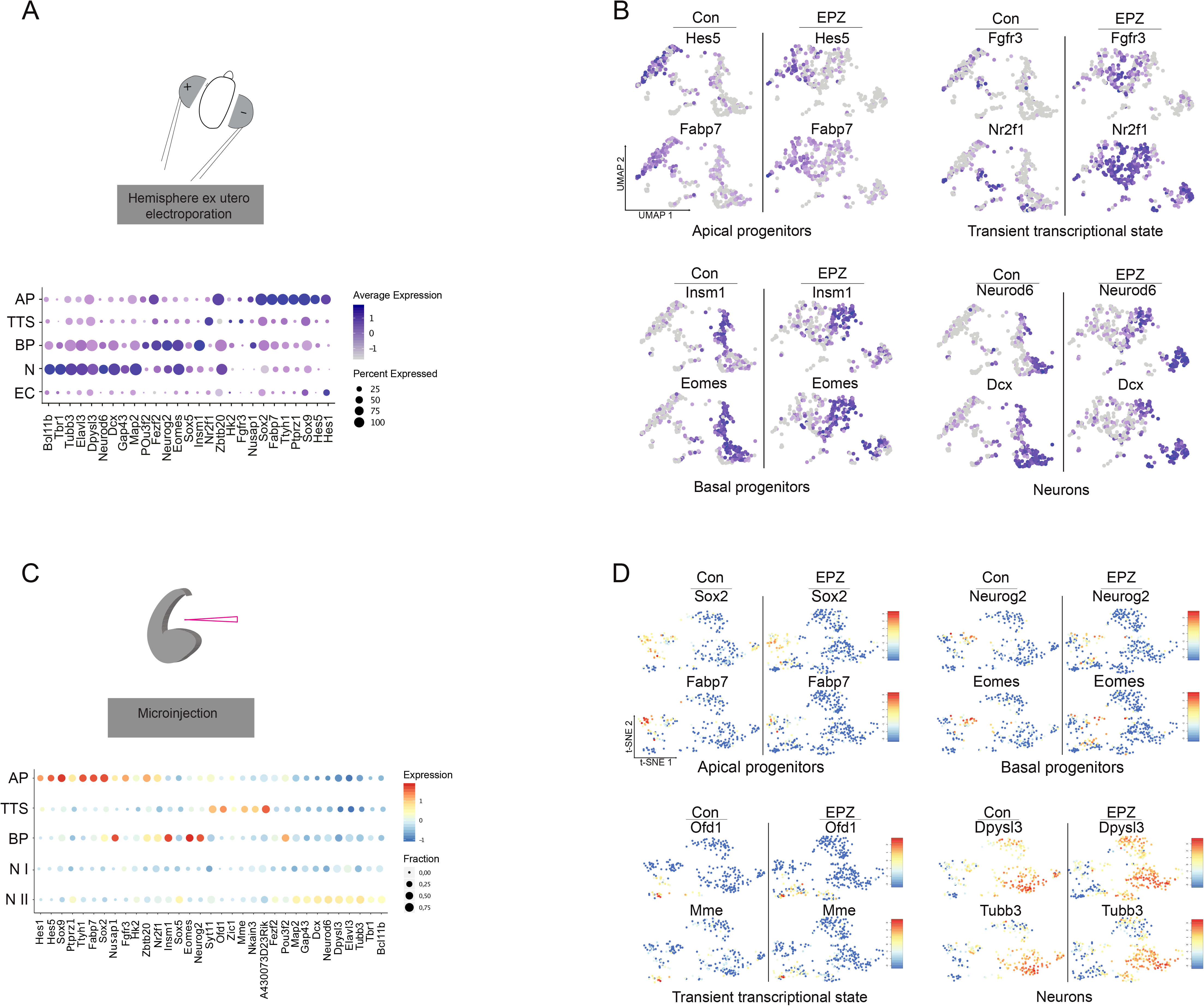
Overview of markers used for cluster annotation. **A)** Dot plot of top markers used in annotation of clusters retrieved from electroporation dataset. **B)** Overview of marker expression (electroporation dataset) in UMAP representation. **C)** Dot plot of top markers used in annotation of clusters retrieved from microinjection dataset. **D)** Overview of marker expression (microinjection data set) in t-SNE representation.

**Supplementary Figure 5.**
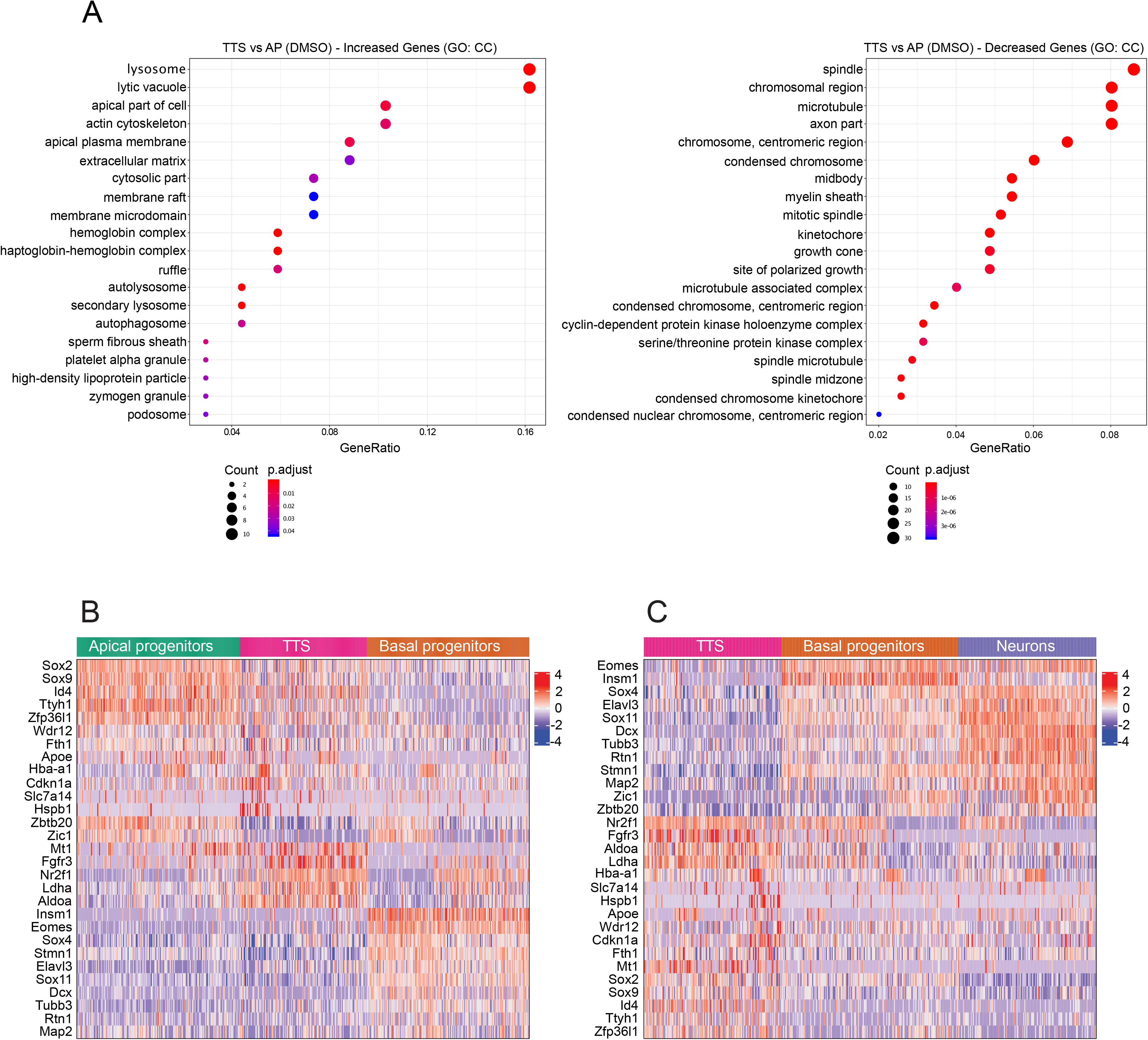
Transcriptional signature of the transient transcriptional state (TTS) which enriches after DOT1L inhibition in electroporated data set. **A)** Left: Top GO terms (p.adjust < 0.01) retrieved using genes upregulated in TTS compared to AP in control condition. Right: Top GO terms (p.adjust < 1e-06) retrieved using genes downregulated in TTS compared to AP in control condition. **B)** Heatmap showing expression levels of selected cell type/state markers clustered for progenitor cells only. **C)** Heatmap showing expression levels of selected cell type/state markers clustered for TTS, BPs and neurons. Scales represent z-scores.

**Supplementary Figure 6.**
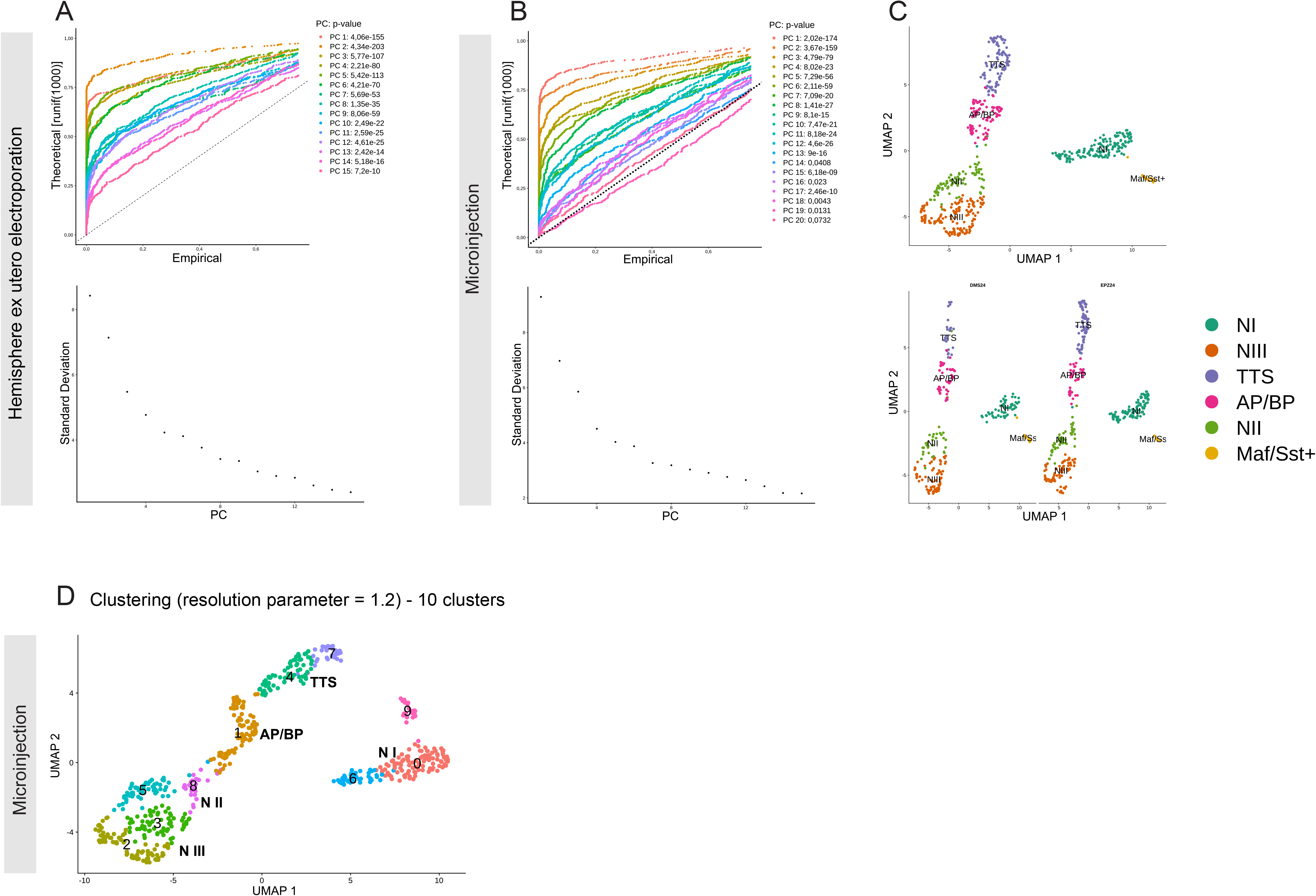
Additional information on parameter optimization for clustering. **A)** Top: JackStraw plot; bottom: Elbow plot from electroporation dataset. **B)** Top: JackStraw plot; bottom: Elbow plot from microinjection dataset. **C)** Clusters retrieved using the top 10 principal components (PC) as input for clustering in microinjection dataset. AP/BP – apical progenitors/basal progenitors, TTS – transient transcriptional state, N I – neurons I, N II – neurons II, N III – neurons III, Maf/Sst+ - Maf positive and Sst positive cluster. **D)** Clusters retrieved using a resolution parameter of 1.2 in the microinjection dataset.

