## supplemental table for "DOT1L activity affects cell lineage progression in the developing brain by controlling metabolic programs"

### Table 1:

**KEY RESOURCES TABLE**

| REAGENT or RESOURCE | SOURCE | IDENTIFIER |
|---|---|---|
| **Primary antibodies** | | |
| Mouse monoclonal anti-dextran antibody, clone DX1 | STEMCELL technologies | Cat# 60026 |
| Rabbit anti-GFP | Abcam | Cat# ab6556 |
| Goat anti-TUBB3 | Everest biotech | Cat# EB11685 |
| Rabbit anti-Tbr2 | Abcam | Cat# ab183991 |
| Mouse anti-g-tubulin | Sigma-Aldrich | Cat# T6557 |
| Mouse anti-TuJ1 | Covance Research Products | Cat# MMS-435P |
| Rabbit anti-Arl13b | Proteintech Group | Cat# 17711-1-AP |
| Rabbit anti-H3K27me3 | Active Motif | Cat# 39155 |
| Rabbit anti-EZH2 | Active Motif | Cat# 39901 |
| **Secondary antibodies** | | |
| Donkey anti-Rabbit IgG (H+L) Highly Cross-Adsorbed Secondary Antibody, Alexa Fluor 488 | Thermo Fisher Scientific | Cat# A-21206 |
| Donkey anti-Rabbit IgG (H+L) Highly Cross-Adsorbed Secondary Antibody, Alexa Fluor 555<br><br>Alexa | Thermo Fisher Scientific | Cat# A-31572 |
| Donkey anti-Mouse IgG (H+L) Highly Cross-Adsorbed Secondary Antibody, Alexa Fluor 488 | Thermo Fisher Scientific | Cat# A-21202 |
| Donkey anti-Mouse IgG (H+L) Highly Cross-Adsorbed Secondary Antibody, Alexa Fluor 647 | Thermo Fisher Scientific Thermo | Cat# A-31571 |
| Donkey anti-Goat IgG (H+L) Cross-Adsorbed Secondary Antibody, Alexa Fluor 647 | Thermo Fisher Scientific | Cat# A-21447 |
| **Tissue culture supplies** | | |
| Rat serum | Charles River Lab., Japan | Cat# S-919-S |

| | | |
|---|---|---|
| DMEM-F12, modified low glucose | Sigma | Cat# D2902 |
| Dextran-Alexafluor 488 (10,000 MW anionic, fixable) | Invitrogen | Cat# D22913 |
| Agarose, low melting point | Roth | Cat# 6351.2 |
| B-27 supplement | Invitrogen | Cat# 17504-044 |
| Cellmatrix type 1 -A | Kyowa chemical products | Cat# 631-00651 |
| L-Glutamine (200 mM) | Thermo fisher scientific | Cat# 25030024 |
| HEPES-NaOH (pH 7.2 1M) | Sigma-Aldrich | Cat# H3537 |
| Slice/hemisphere culture incubation box | MPI-CBG workshop | N/A |
| **Chemicals, peptides** | | |
| EPZ5676 | Selleckchem | Cat# S7062 |
| DMSO | Thermo Fisher Scientific | Cat# TS-20688 |
| L-Albizziine | Gold Biotechnology | Cat# A-230-250 |
| **Commercial assays** | | |
| Neural Tissue Dissociation Kit (P) | Miltenyi Biotec | Cat# 130-092-628 |
| **Deposited data** | | |
| Single cell RNA sequencing (scRNA-seq) data | This paper | GEO: GSE176323 |
| Experimental models: Organisms/strains | | |
| Wild type mouse: C57BL/6JRj | Janvier Labs | N/A |
| Transgenic mouse: Tis21+/tm2(Gfp)Wbh | (Haubensak et al., 2004) | N/A |
| **Plasmids and Oligonucleotides** | | |

| | | |
|---|---|---|
| pCAGGS-mCherry | Genscript | N/A |
| pCMV-ASNS-GFP | Origene | MG208883 |
| pCMV-GFP | Origene | PS100010 |
| 192 polyT primers with unique molecular index and cell barcode | Integrated DNA technologies | Sagar et al., 2018 |
| CEBPG_BS1_ASNS_fw: CAGCATGGATTTGCAGTCAT | Sigma | N/A |
| CEBPG_BS1_ASNS_rv: TTTCAGGTGGGAGAGAGGATT | Sigma | N/A |
| CEBPG_BS2_ASNS_fw: AAGCCAGGGTCATCTAGGAAG | Sigma | N/A |
| CEBPG_BS2_ASNS_rv: GCAGAGACATCCCACATCAAT | Sigma | N/A |
| ASNS_TSS_fw: TCCCGCTTACCTGAGCACTA | Sigma | N/A |
| ASNS_TSS_rv: CAGCCACATGATGAAACTTCC | Sigma | N/A |
| randomhexRT primer, GCCTTGGCACCCGAGAATTCCANNNNNN | Integrated DNA technologies | Sagar et al., 2018 |
| RNA PCR Primers, sequences available from Illumina (RP1, RPI1-RPI12, TruSeq) | Integrated DNA technologies | Sagar et al., 2018 |
| **Tissue and hemisphere culture supplies** | | |
| 40% $O_2$, 55% $N_2$, 5% $CO_2$ | Airliquide | N/A |
| Sodium hydroxide pellets | Merck | Cat# 106482 |
| Tyrode's salt | Sigma-Aldrich | Cat# T2145-10x1L |
| Sodium bicarbonate | Merck | Cat# 106323 |
| Penicillin-Streptomycin | Gentaur | Cat# PAA P11-010 |
| **Software and algorithms** | | |
| RaceID3 | v0.1.6 | https://github.com/dgrun/RaceID3_StemID2_package |

| | | |
|---|---|---|
| RStudio | v.3.6.1 | https://www.rstudio.com/ |
| R | v1.2.5042 | https://www.r-project.org/ |
| Seurat | v.3.1.2 | https://satijalab.org/seurat/ |
| Scran | v3.12 | https://bioconductor.org/packages/release/bioc/html/scran.html |
| SingleCellExperiment | v1.8.0 | https://www.bioconductor.org/packages/release/bioc/html/SingleCellExperiment.html |
| SCENIC | v1.1.2 | https://github.com/aertslab/SCENIC |
| GENIE3 | v1.12.0 | https://github.com/aertslab/GENIE3 |
| RcisTarget | v1.10.0 | https://www.bioconductor.org/packages/release/bioc/html/RcisTarget.html |
| AUCell | v1.12.0 | https://bioconductor.org/packages/release/bioc/html/AUCell.html |
| Bioconductor | v3.12 | https://www.bioconductor.org/ |
| clusterProfiler | v3.14.0 | https://bioconductor.org/packages/release/bioc/html/clusterProfiler.html |
| Prism | v.8.0. | https://www.graphpad.com/scientific-software/prism/ |
| **Single cell RNA sequencing reagents and supplies** | | |

| | | |
|---|---|---|
| RNaseOUT | Invitrogen | Cat# 10777-019 |
| Superscript II | Invitrogen | Cat# 18064-014 |
| Second Strand Buffer | Invitrogen | Cat# 10812-014 |
| E. coli DNA ligase | Invitrogen | Cat# 18052-019 |
| E. coli RNaseH | Invitrogen | Cat# 18021-071 |
| E. coli DNA polymerase | Invitrogen | Cat# 18010-025 |
| AMPure XP beads | Beckman Coulter | Cat# A63880 |
| RNAClean XP beads | Beckman Coulter | Cat# A63987 |
| MEGAscript T7 Transcription Kit | Invitrogen | Cat# AM1334 |
| Phusion High-Fidelity PCR Master Mix with HF Buffer | NEB | Cat# M0531 |
| ExoSAP-IT For PCR Product Clean-Up | Affymetrix | Cat# 78200 |
| NEBNext Magnesium RNA Fragmentation Module | NEB | Cat# E6150S |
| randomhexRT primer, GCCTTGGCACCCGAGAATTCCANNNNNN | Integrated DNA technologies | Sagar et al., 2018 |
| RNA PCR Primers, sequences available from Illumina (RP1, RPI1-RPI12, TruSeq) | Integrated DNA technologies | Sagar et al., 2018 |
| 192 polyT primers with unique molecular index and cell barcode | Integrated DNA technologies | Sagar et al., 2018 |
| **Other** | | |
| scRNA-seq data analyses workflows | This paper | https://github.com/Vogel-lab/DOT1L_activity_neocortex-paper |
